# A sequence variation between two orthologues alters functional expression of the potassium channel Kesv of *Ectocarpus siliculosus* virus

**DOI:** 10.64898/2026.07.31.740523

**Authors:** Purva Asrani, Aladdin Elgendy, André P. Zeipelt, Gunnar Goerges, Julian A. Schreiber, Richard Brown, Daniel Todt, Daniel Tapken, Lars V. Schäfer, Guiscard Seebohm, Raphael Stoll

**Affiliations:** Biomolecular Spectroscopy and RUBiospec, NMR, Faculty of Chemistry and Biochemistry, Ruhr University of Bochum, D-44801, Bochum, Germany; Center for Theoretical Chemistry, Faculty of Chemistry and Biochemistry, Ruhr University of Bochum, D-44801, Bochum, Germany; Institute for Genetics of Heart Diseases (IfGH), Department of Cardiovascular Medicine, University Hospital Münster, D-48149 Münster, Germany; Institute for Pharmaceutical and Medicinal Chemistry, Westfälische Wilhelms-Universität Münster, Corrensstraße 48, D-48149 Münster, Germany; Department of Translational and Computational Infection Research (TRACiR), Medical Faculty, Ruhr University of Bochum, D-44801, Bochum, Germany; European Virus Bioinformatic Center (EVBC), Jena, Germany; Receptor Biochemistry, Faculty of Chemistry and Biochemistry, Ruhr University of Bochum, D-44801, Bochum, Germany

**Keywords:** Host-virus interactions, Kesv, *Ectocarpus siliculosus*, *Xenopus laevis*, Potassium ion channel, voltage clamp

## Abstract

The potassium channel Kesv encoded by the *Ectocarpus siliculosus* virus (Kesv 1) differs by seven amino acid residues from its host-derived homolog (Kesv 2), resulting from lysogenic integration. When expressed in *Xenopus laevis* oocytes, Kesv 1 displayed significantly higher ion conductance and functional expression than Kesv 2, as demonstrated by GFP fluorescence and voltage clamp measurements. This study provides the first structural and functional analysis of Kesv 2, uncovering key differences between the original and host-derived variant. The systematic residue substitutions – based on location– from Kesv 2 to the corresponding residues in Kesv 1 illustrated that two amino acid exchanges in close proximity to the pore region (Q61H and T66A), albeit not individually but in combination, significantly resulted in a loss-of-function phenotype in Kesv 1. AlphaFold predictions and subsequent molecular dynamics simulations did not reveal significant differences between Kesv 1 and Kesv 2 structural models, suggesting that the loss of function cannot be attributed to differences at the structural level. Instead, a reduced surface expression of Kesv 2, caused by the sequence modulations in the brown algal host, appears more plausible. Notably, the pharmacological profiling with Linopirdine and Sotalol highlights differences in drug sensitivity, establishing these minimalist channels (core channel structure without regulatory domains) as tractable models for dissecting novel fundamental principles of ion channel function and drug interaction, while highlighting key differences from more complex channel systems.

**Significance Statement:** Potassium channels are essential for cellular excitability, yet their large size and structural complexity limit our understanding of the core features underlying channel function. Here, we identified and established an orthologous model to compare the effects of evolutionarily acquired mutations in two voltage-sensing potassium channels-the viral potassium channel from Ectocarpus siliculosus virus (Kesv 1) and its host-homolog derivative (Kesv 2) as simplified model systems for understanding ion channel physiology and host-viral interactions. We provide the first functional characterization of Kesv 2 in Xenopus laevis oocytes using two-electrode voltage-clamp and site-directed mutagenesis, revealing that, despite sharing an identical SVGYG selectivity-filter motif and differing by only 7 residues, they exhibit distinct ion-conduction properties.

## 1. Introduction

Host-virus interactions often serve in exploring the basic to complex biological mechanisms between the two underlying species (1). Such understanding is based on the fact that viruses hijack the cellular machinery of the host cell to undergo transcription, translation, and other biochemical pathways to facilitate their propagation and survival (2). Viruses undergoing lysogenic life cycles can stably insert their genome into the host organisms and can remain dormant for long periods of time (3, 4). On the one hand, this phenomenon serves as an excellent model system to understand evolutionary developmental processes. On the other hand, the minimalist structure of viral proteins is used to understand the architectural designs of more complex proteins homologous to those in higher organisms. So far, many independent studies on host-virus interactions have been conducted. However, the structural-functional changes in viral proteins associated with evolutionarily driven mutations have barely been explored.

An interesting aspect of studying host-virus interactions is to explore how hosts defend themselves against viral infections. One way is to introduce mutations that reduce the expression of critical genes required for viral replication, while other mechanisms involve the suppression of expression at the transcriptional level (5, 6). Along the lines of these concepts, we focused on understanding the host-virus interaction between *Ectocarpus siliculosus*, a marine brown alga and a lysogenic virus, *Ectocarpus siliculosus* virus (EsV-1). *Ectocarpus siliculosus* has a unique evolutionary lineage. It has evolved independently of plants, animals, and fungi as it exhibits a complex multicellularity, which also renders it a prime model for understanding evolutionary developmental processes (7). However, under natural conditions, EsV-1 infections are very prevalent across the species of *Ectocarpus* and mainly infecting through a lysogenic process (8). EsV-1 is a large dsDNA virus with a 335 kbp genome comprising 231 genes (9). One of the several crucial proteins of this virus is Kesv, a potassium ion channel that may play an important role in initiating the host-virus interaction. This small homotetrameric protein of 124 amino acids per monomer forms a functional K^+^ selective ion channel, characterized by a stable ion-binding complex within the selectivity filter (SF), upon membrane localization (10). In this study, we focus on understanding the effects of evolutionarily acquired mutations between two orthologs expressed as voltage-sensing potassium channels, referred to as Kesv 1 and Kesv 2. Despite sharing an identical SVGYG selectivity filter motif and differing by only 7 residues, they exhibit distinct ion-conduction properties. To our knowledge, this study presents the first comprehensive functional characterization of Kesv 2, revealing how minimal sequence variations can critically modulate viral channel expression.

Like other K^+^ channels (11), Kesv forms a tetrameric assembly in the membrane. Each monomer harbors α-helical transmembrane segments: M0 (N-terminal) helix, M1 helix, Pore (P) helix and M2 helix (12). The helices M1 and M2 are connected via an extracellular loop and within the pore domain, which harbors a short helical segment (P helix) and the signature SF motif (SVGYG sequence), also known as the pore loop. The pore domain is the core of a channel that enables ion permeation, with K^+^ specificity determined by the SF. The ion-binding sites of the SF are formed by the amino acid backbone carbonyl groups, providing an ideal coordination geometry for K^+^ ions. In this way, there are five binding sites, S0–S4, with S0 located on the extracellular side and S4 on the intracellular side (13). Any mutations in these regions usually significantly alter the ion channel’s activity and, hence, are deemed critical for channel function. Interestingly, the pore of K^+^ channels is conserved across all species, whereas variations are predominant within the regulatory domains, which often are generally species-specific (14). These regulatory domains may confer additional properties to the channel, including the sensing of external stimuli or interactions with different ligands for signaling (14). Kesv lacks regulatory domains yet forms a complete ion channel; hence, it can be considered minimal in its architecture. Even with atomic-level insights into its structure, the way in which the structure governs the protein’s function remains poorly understood. Both the smaller size and the homology of the pore domain of Kesv to the more complex eukaryotic ion channel proteins, like Kv channels, make these channels interesting targets for studying their structure and function, and for understanding of pharmacological mechanisms (15).

The majority of metabolic, neurodegenerative and cardiac diseases in humans are linked to the abnormalities in the gating mechanism that restrict the opening or closing of K^+^ channels (16, 17). Given the growing incidence of diseases such as epilepsy, long QT syndrome, and type 2 diabetes mellitus linked to channel proteins (18), studying the interactions of potassium channels with other molecules may help facilitate the development of targeted therapeutic strategies (19). Therefore, the search for agonists and antagonists in these cases is central to drug screening processes. In order to investigate the pharmacological potential of various modulators, the underlying fundamental principles of ion channel physiology and drug interactions need to be explored. The availability of high-resolution structures may facilitate such understanding of the behavior of these molecules (20–22). However, billions of protein sequences exist in databases, far from being purified (23) thereby limiting our ability to obtain functional and structural insights. AlphaFold has enabled computational prediction of protein structures and has been successfully used in many applications (24). However, when used as a tool for generating structural hypotheses, AlphaFold predictions should be interpreted with caution and experimentally validated whenever possible.

## Results

### Kesv 1 is an active ion channel with higher functional expression than Kesv 2

The alignment of the Kesv 1 and Kesv 2 sequences (both 124 amino acids long) shows seven distinct amino acid substitutions (7/124) distributed across different regions of the ion channel (Fig. 1A). Kesv 1 is identical to the potassium ion channel protein of EsV-1 (10), whereas Kesv 2 represents the integrated version of Kesv 1 within the genome of *Ectocarpus siliculosus* brown algae (25). The first two gene variants, A7V and V20A, are in the predicted M0 helix of the channel. The gene variants, Q61H and T66A occur towards the end of the predicted M1 helix (near the pore helix), while the gene variant L59M lies within the M1 helix. The two C-terminal gene variants, T103V and A119V, are positioned in the predicted M2 helix.

**Figure 1.**
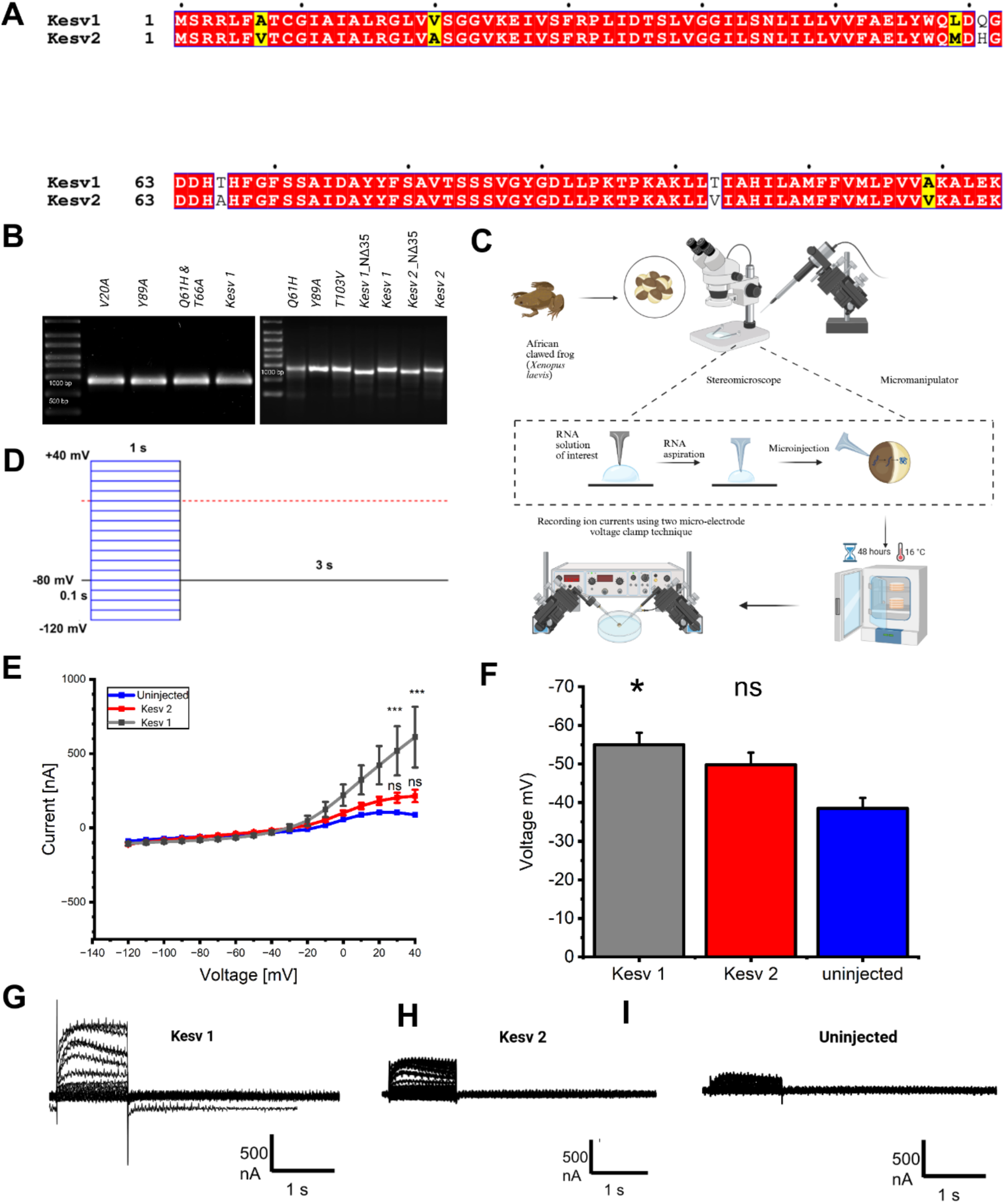
Functional expression of Kesv 1 and Kesv 2 potassium channel in *Xenopus laevis* oocytes. (A) Sequence alignment of Kesv 1 and Kesv 2. The red to white highlighting scheme is a measure of similarity between residues, with red being equivalent, yellow being similar and white being dissimilar. (B) Agarose gel electrophoresis showing *in vitro* transcription products of Kesv 1 (wild type), mutagenised Kesv variants (systematic substitution of Kesv 2 to Kesv 1), and N-terminal truncated variants. (C) Oocytes obtained from the African clawed frog (*Xenopus laevis*) are selected under a stereomicroscope and positioned using a micromanipulator. RNA of interest is aspirated into a microinjection pipette and injected into defolliculated oocytes. Injected oocytes are incubated at 16 °C for approximately 48 hours to allow heterologous expression of the encoded ion channels. Functional expressions are assessed by recording ionic currents using the TEVC technique, in which membrane potential is controlled and resulting currents are measured using two intracellular microelectrodes. (D) Schematic of the voltage-clamp protocol used for electrophysiological recordings. Cells were held at −80 mV and stepped from −120 mV to +40 mV for 1 s, with 3 s intervals at the holding potential between sweeps. (E) I-V curve of oocytes expressing Kesv 1 (gray; n=4), Kesv 2 (red; n=5), and uninjected oocytes (blue; n=4). (F) Bar graph showing the resting membrane potential of oocytes shown in panel E. (G–I) Representative current traces recorded in response to voltage steps from −120 mV to +40 mV in (G) Kesv-1, (H) Kesv-2, and (I) uninjected oocytes. Statistical significance was determined using one-way ANOVA followed by Tukey’s post hoc test. ns, *p* > 0.05; *p* < 0.05; \**p* < 0.01; \*\**p* < 0.001. Created in https://BioRender.com

The functional expression of *in vitro* transcribed Kesv 1 and Kesv 2 was examined electrophysiologically in *Xenopus laevis* oocytes using two electrode voltage clamp technique TEVC (26) (Fig. 1 B, C). The comparisons were based on differential ion conductance between uninjected oocytes and oocytes injected with cRNA provided with one-second test pulse across the voltage range of −120 to +40 mV (Fig. 1D). Expression of Kesv 1 resulted in significantly increased outward currents from +30 to +40 mV compared to uninjected oocytes. Over the complete voltage range (−120 to +40 mV), currents recorded for Kesv 2 were substantially lower than those recorded for Kesv 1 and were closer to those of uninjected oocytes (SI Table S3a). For example, at 50 mM external K^+^ concentration and +40 mV, the current observed for Kesv 1 was 611 ± 205 nA (n=4) and for Kesv 2 was 215 ± 42 nA (n=5). Uninjected, in comparison, only showed currents of 87 ± 9 nA (n=4) at +40 mV (Fig. 1E, G-I).

Similarly, the resting potentials were compared for both constructs (Fig. 1F). Compared to uninjected oocytes (−39 ± 3 mV), oocytes expressing Kesv 1 showed a significantly lower resting potential (−55 ± 3 mV) at 96 mM K^+^ external concentration. In contrast, Kesv 2-expressing oocytes showed a less negative resting potential (−50 ± 3 mV) than Kesv 1-expressing ones (SI Table S3b). The results suggest that Kesv 2 is impaired but still functional enough to maintain a resting potential comparable to Kesv 1.

The highly significant current differences and reduced resting potential observed for Kesv 1 in comparison to tiny currents of uninjected oocytes suggested functional expression of Kesv 1 as a functional potassium channel. Moreover, Kesv 2 also elicited currents but not significantly different to uninjected. Hence, their reduced functional expression suggested the possible effect of those seven gene variants in either reducing the ion conductance properties or the surface expression of the potassium ion channel. Therefore, further investigations to further understanding of impact of these gene variants on structure, localization or functional change of ion channels were carried out, as discussed in the following sections.

#### Effect of mutations in M0 helix

The currents were recorded by standard TEVC measurements (24) when oocytes injected with cRNA of Kesv 1 (wild type), V20A (test mutant), Y89A (negative control) and uninjected oocytes were elicited with one-second test pulse across the voltage range of −120 to +40 mV at a holding potential of −80 mV (Fig. 2A).

**Figure 2.**
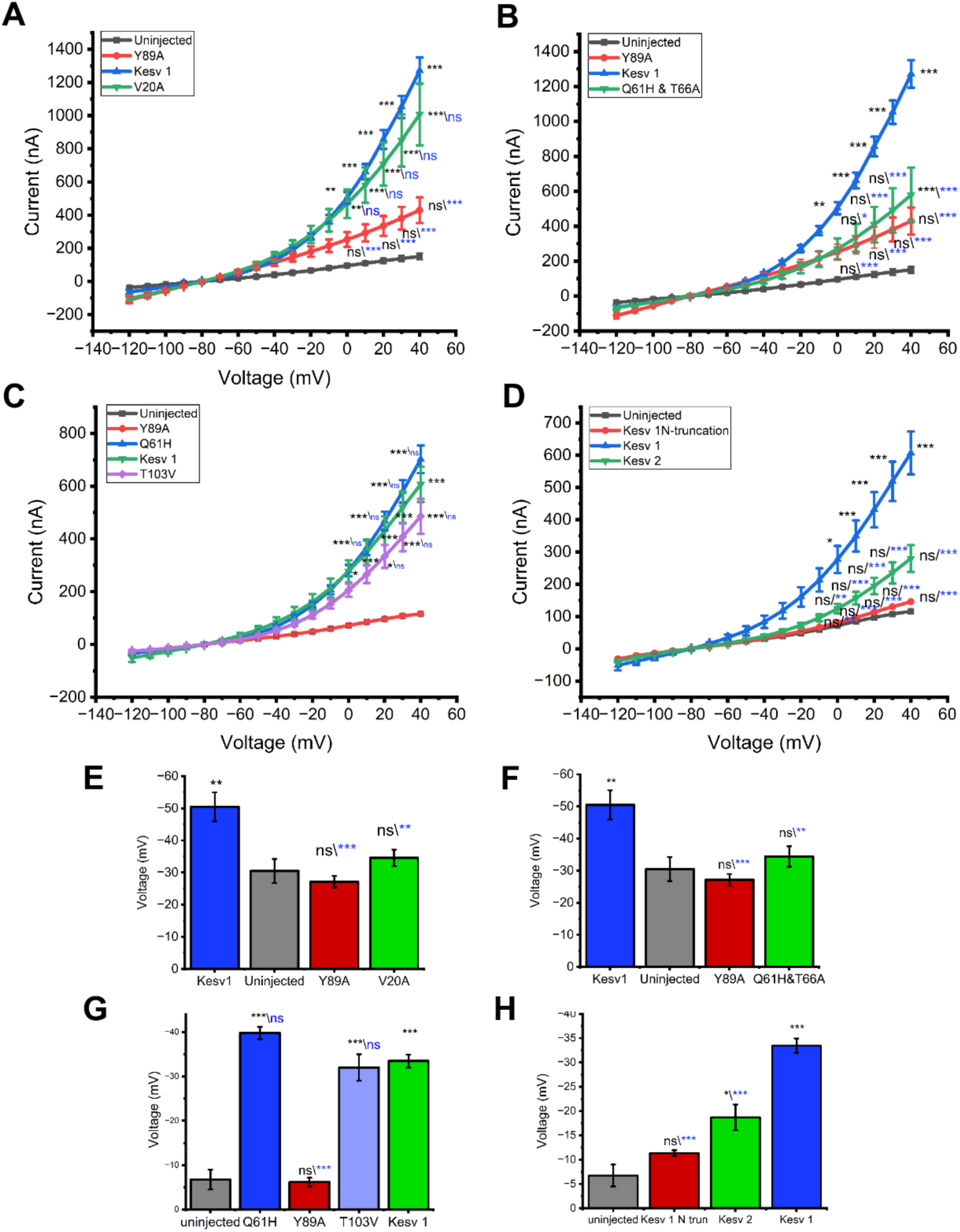
Functional expression of Kesv-1 and its mutants in *Xenopus* oocytes based on their structural locations. (A) Effect of M0 helix mutations on I–V curves of oocytes expressing Kesv-1 (blue, *n* = 12), V20A (green, *n* = 12), Y89A (red, *n* = 12), and uninjected oocytes (gray, *n* = 8). (B) Effect of mutations near the pore helix on I–V curves of oocytes expressing Kesv-1 (blue, *n* = 12), Q61H/T66A (green, *n* = 10), Y89A (red, *n* = 12), and uninjected oocytes (gray, *n* = 8). (C) Effect of M1 and M2 helix mutations on I–V curves of oocytes expressing Kesv-1 (green, *n* = 15), Q61H (blue, *n* = 10), T103V (purple, *n* = 7), Y89A (red, *n* = 5), and uninjected oocytes (gray, *n* = 4). (D) Effect of N-terminal/M0 helix truncation on I–V curves of oocytes expressing Kesv-1 (blue, *n* = 15), Kesv-2 (green, *n* = 13), Kesv-1-NΔ35 (red, *n* = 6), and uninjected oocytes (gray, *n* = 4). (E–H) Bar graphs showing the resting membrane potential of the corresponding oocytes shown in panels (A–D), respectively. Statistical significance was determined using one-way ANOVA followed by Tukey’s post hoc test. ns, *p* > 0.05; *p* < 0.05; \**p* < 0.01; \*\**p* < 0.001.

The whole cell currents observed at 50 mM external concentration of K^+^ ions at +40 mV averaged at 1272 ± 79 nA for Kesv 1 (n=12); 1007 ± 187 nA for V20A (n=12); 430 ± 77 nA for Y89A (n=12) and 150 ± 20 nA for uninjected oocytes (n=8) (Fig. 2A). The functional expression of V20A was highly significant from +10 to +40 mV in comparison to the uninjected oocytes, suggesting a functional expression however not significantly different to Kesv 1 (across all voltages) suggesting that residue V20A does not impact plasma membrane expression of the channel protein. Furthermore, the I-V plot also confirms Kesv 1 as a functionally active ion channel exhibiting highly significant currents from 0 to +40 mV, on the contrary, Y89A currents were only slightly larger and not significantly different to uninjected (SI Table S4a). Moreover, currents of Y89A significantly differ from Kesv 1 (across all voltages), proving the loss of ion channel activity upon mutations in the selectivity filter (10).

The comparison, based on resting potential (Fig. 2E) showed an opposite trend where Kesv 1 (−51 ± 5 mV) did show a significant drop in resting potential in comparison to uninjected (−30.5 ± 4 mV); however, V20A (−35 ± 3 mV) remained non-significant to uninjected and significantly different to Kesv 1. This implies a possible role of V20A in reducing the functionality of Kesv 1 at least to some extent, if not a major contributor to a functional change, in contrast to the observation based on the I-V plot (SI Table S4b). Consistently, Y89A (−27 ± 2 mV), showed resting potentials similar to those of uninjected oocytes.

#### Effect of mutations near the pore helix

The double mutant Q61H & T66A (test mutant) was injected into oocytes alongside Y89A (negative control) and Kesv 1 (wild type). All injected and uninjected oocytes were elicited with one-second test pulse across voltage range of −120 to +40 mV.

The average currents for Q61H and T66A (579 ± 155 nA; n=10) were significantly lower in comparison to Kesv 1 (1272 ± 79 nA; n=12) (across all voltages) and significantly different to uninjected (151 ± 20 nA; n=8) only at +40 mV (Fig. 2B). Moreover, Kesv 1 exhibited the highest currents significantly different to uninjected at +0 to +40 mV while Y89A (n =12) remained non-significant to uninjected with an average current of 430 ±77 nA (SI Table S5a). Of note, the Q61H mutation increased the ion conductance properties of Kesv 1 *(section: Effect of mutations in M1 and M2 helix),* however; in combination with T66A, it was shown to reduce the conductivity significantly. These findings prove the double mutant to be the major contributor to the functional change and is responsible for loss of function mutation of Kesv 1.

Further comparisons on the basis of resting potential (Fig. 2F) also followed a similar trend, where Q61H & T66A (−34 ± 3 mV) did not result in a significant drop in resting potential as compared to uninjected oocytes (−31 ± 4 mV); however, they remained significantly different to Kesv 1 (−51 ± 5 mV). Y89A, on the other hand, exhibited a resting potential of −27 ± 2 mV, non-significant to uninjected and significantly different to Kesv 1 (SI Table S5b).

#### Effect of mutations in M1 and M2 helix

The amino acid exchanges in the M1 helix (Q61H) and M2 helix (T103V) were chosen as test mutants to understand their impact on the functional expression of Kesv 1 (Fig. 2C). Uninjected and oocytes injected with cRNA of T103V (test mutant 1), Q61H (test mutant 2), Kesv 1 (wild type) and Y89A (negative control) were subjected to standard TEVC measurement (24) for recording the ion conductance through one-second test pulses across the voltage range of −120 to + 40 mV.

The average currents obtained for Q61H (702 ± 52 nA; n=10) was significantly higher from +10 to +40 mV than for the uninjected (116 ± 11 nA; n=4) and slightly higher than Kesv 1 (607 ± 67 nA; n=15) but not significantly. T103V, on the other hand, showed highly significant currents (486 ± 66 nA; n=7) in comparison to uninjected and non-significant to Kesv 1 (across all voltages), suggesting less impact of both mutations on functional level and hence not the major contributors to the change of channel activity. Kesv 1 again showed highly significant results in comparison to uninjected across the voltage range from 0 to + 40 mV however, Y89A currents (116 ± 5 nA; n=5) were similar to those of uninjected, while being significantly different to Kesv 1 (SI Table S6a) (Fig. 2C).

Observations based on a drop in resting potential are consistent with the I-V curve, indicating no role of these mutants on the functionality of an ion channel (Fig. 2G). The resting potential of the two mutants, Q61H (−40 ±1 mV) and T103V (−32 ± 3 mV), was significantly different to uninjected (−6.75 ±2.25 mV) but not significantly different to Kesv 1 (−33 ± 1 mV). Kesv 1, on the other hand, continued to exhibit a statistically significant drop in resting potential in comparison to uninjected, while Y89A (−6 ± 1 mV) showed similar resting potentials (SI Table S6b).

#### Effect of M0 helix truncation

The first 35 residues of the N-terminus (including the M0 helix and the loop region) were removed from Kesv 1 (Kesv 1-NΔ35). The truncated Kesv 1-NΔ35 (test mutant) was injected into oocytes alongside Kesv 1 (wild type) and Kesv 2 (mutant), and similar TEVC measurements were performed as described in the above sections (Fig. 2D).

Kesv 1-NΔ35 showed highly reduced functional expression with an average current of 146 ± 5 nA (n=6) in comparison to Kesv 1 (607 ± 67 nA, n=15) and remained non-significantly different to uninjected (116 ± 11 nA; n=4) across all voltages tested. Additionally, Kesv 1 showed significantly higher expression than uninjected from +0 to +40 mV, whereas Kesv 2 (280 ± 41 nA; n=13) remained non-significantly different to uninjected but significantly different to Kesv 1 across all voltages (SI Table S7a).

Resting potential comparisons showed a highly significant drop for Kesv 1 (−33 ± 1 mV) in comparison to uninjected (−7 ± 2 mV) but no significant differences for the Kesv 1-NΔ35 (−11 ± 1 mV) were observed (Fig. 2H). It also remained significantly different to Kesv 1, supporting the notion that the *N-terminal ⍺-helix* is an essential component of this channel protein, perhaps being required for proper interaction with and/or insertion into the membrane. However, Kesv 2 (−19 ± 3 mV), in this case, exhibited a significantly different resting potential compared to uninjected but nevertheless was still significantly different to Kesv 1 (SI Table S7b).

### Pharmacological characterization reveals noncanonical modulation of Kesv

The effect of pharmacological agents on the channel activity was determined by continuous perfusion of compounds on oocytes expressing Kesv 1 using standard TEVC measurements (26) (Fig.3 A-D).

**Figure 3.**
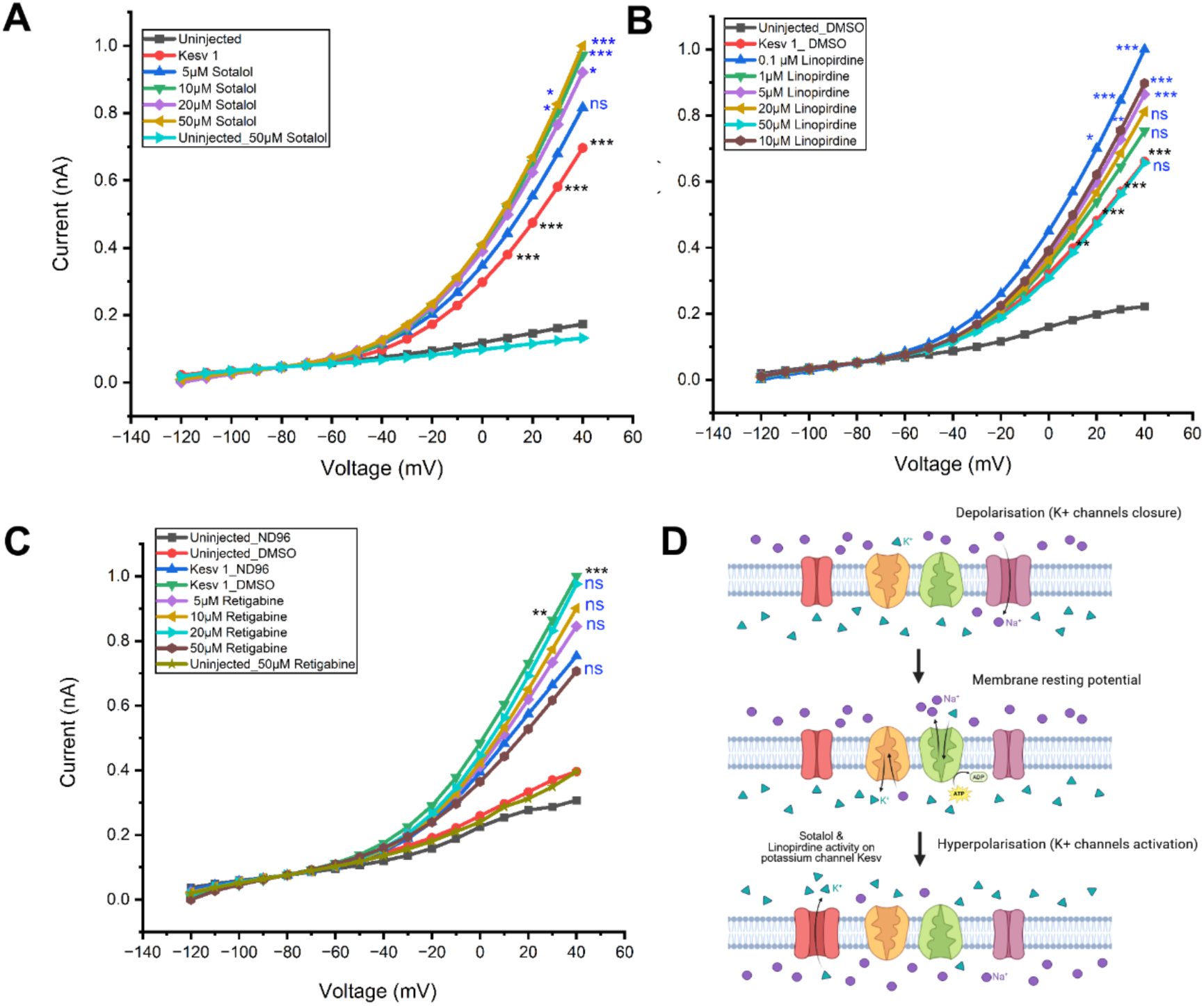
Pharmacological characterization of Kesv channel function. (A-C) The pharmacological effect of drugs (A) Sotalol (B) Linopirdine and (C) Retigabine on functional expression of Kesv 1 protein. Statistical significance is determined using one-way ANOVA followed by Tukey’s post hoc test. ns, *p* > 0.05; *p* < 0.05; \**p* < 0.01; \*\**p* < 0.001. All currents are normalized to I_max_. (D) Schematic representation of Kesv channel modulation. Membrane depolarization (via Na⁺ channel activation) triggers opening of K⁺ channels, leading to K⁺ efflux and restoration of the resting membrane potential. Pharmacological activation of Kesv by Sotalol and Linopirdine enhances channel activity, promoting a shift toward a hyperpolarized membrane state. Created in https://BioRender.com

#### Sotalol

Sotalol at four different concentrations, i.e., 5 μM, 10 μM, 20 μM and 50 μM, was perfused on oocytes expressing Kesv 1 channel protein, and in general, increase in ion conductance was detected (Fig. 3A). The observed currents at each concentration and voltage were statistically compared to the currents recorded for functional expression of Kesv 1 to its corresponding voltage. The highest currents were observed at 50 μM Sotalol concentration. The currents were significant at +30 mV (*, p < 0.05) and (***, p < 0.001) at +40 mV. When examining other concentrations, such as 5 μM, no significant increase in currents were reported across any tested voltage while, a concentration of 20 μM, slightly performed better by reporting an increase at +40 mV, significantly different to Kesv 1 wild type (*, p < 0.05). The 10 μM concentration also showed significantly increased ion conductance at +30 mV (*, p < 0.05) and + 40 mV (***, p < 0.001) than Kesv 1, a pattern like 50 μM concentration, appearing only slightly lower in its current amplitude (SI Table S8). Therefore, Sotalol appeared to be a weak activator of Kesv 1.

The oocytes expressing Kesv 1 were also highly significantly different as compared to uninjected oocytes, across voltages starting from + 10 to +40 mV (***, p < 0.001), indicating the functional expression of channel protein. Uninjected oocytes treated with the highest concentration of compound, i.e. 50 μM Sotalol, did not show any significant changes and exhibited a slight reduction in expression of endogenous channel proteins.

#### Linopirdine

The activity of Kesv 1 was recorded at six different concentrations of Linopirdine, i.e. 0.1 μM, 1 μM, 5 μM, 10 μM, 20 μM and 50 μM. Figure 3B shows an I-V graph highlighting the effect of Linopirdine on Kesv 1-expressing oocytes using standard TEVC measurements (26). Linopirdine was also found to be an activator of Kesv 1 channel, where even very small concentration of 0.1 μM, was found capable of increasing currents to a highly significant level (Fig. 3B). The currents were significantly different at + 20 mV (*, p < 0.05) and highly significantly different at + 30 (***, p < 0.001) and + 40 mV (***, p < 0.001). Linopirdine thus exhibited a strong activating effect on Kesv 1. Following this, the second highest increased currents, significantly different at +30 mV (*, p < 0.05) and highly significantly different at + 40 mV (***, p < 0.001) were observed at a concentration of 10 μM. Similarly, 5 μM at + 40 mV (***, p < 0.001) produced significantly different increased currents in comparison to Kesv 1 wild type expression. Other concentrations like 1 μM, 20 μM and 50 μM, however, did not result in any significant change (ns). Significant differences were observed when Kesv 1-injected oocytes were compared with uninjected oocytes under 0.5% DMSO as a control. The currents were found to be significantly different at voltage of +20 to +40 mV (***, p < 0.001) (SI Table S9).

#### Retigabine

Likewise, Sotalol, four different concentrations of Retigabine, i.e., 5 μM, 10 μM, 20 μM and 50 μM, were studied by continuous perfusion of compound on Kesv 1 expressing oocytes under standard TEVC measurements (26). The figure 3C shows an I-V graph of the effect of Retigabine on oocytes expressing Kesv 1 channel protein. While Kesv 1-DMSO did show statistically significant differences to uninjected-DMSO oocytes at voltages from + 30 mV (**, p < 0.01) and +40 mV (**, p < 0.001), the effect of compound on Kesv 1 was not significant at any concentration and any tested voltage (SI Table S10). The highest current remained for Kesv 1, and all other concentrations followed next but did not show any statistically significant results, highlighting the lack of Retigabine’s effect on Kesv 1 channel (Fig. 3C).

#### GFP fluorescence imaging shows reduced surface expression of Kesv 2

The analysis of GFP fluorescence expression in oocytes expressing all three GFP-fusion constructs, Kesv 1– GFP, Kesv 2 - GFP, and Kesv 1 serving as a control, were imaged using an epifluorescence microscope (Fig. 4 A-C). Imaging conducted at 40% excitation (Fig. 4B) used a minimum exposure time of 1/1.5 s whereas 0.25 s of minimum exposure was recorded for constructs under 100% excitation intensity (Fig. 4C).

**Figure 4.**
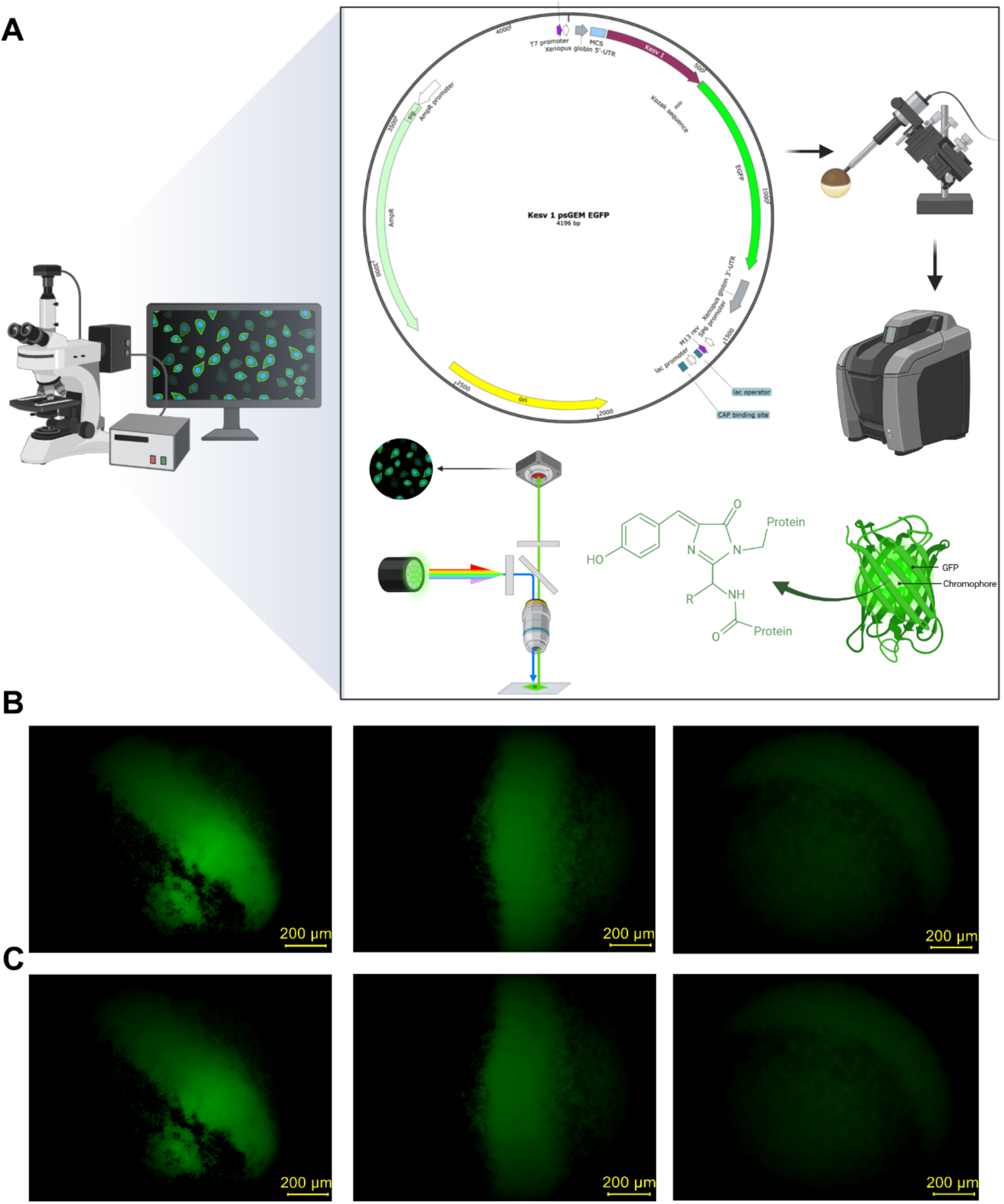
Fluorescence-based expression analysis of Kesv variants in *Xenopus* oocytes (A) Schematic representation of Kesv 1–EGFP fluorescence expression. Kesv 1 was cloned into the psGEM-EGFP vector and expressed in oocytes following microinjection. Fluorescence imaging was performed using an epifluorescence microscope, where excitation light was directed onto the sample through a dichroic mirror and the emitted GFP fluorescence was collected by the detector. The GFP chromophore emits fluorescence at a longer wavelength upon excitation, enabling visualization of Kesv 1 expression in oocytes. (B-C) Fluorescence microscopy images of Xenopus *laevis* oocytes expressing Kesv 1-EGFP (left), Kesv 2-EGFP (middle) and oocytes expressing Kesv 1 without EGFP tag-plasmid (right) at (B) 40% excitation intensity and (C) 100% excitation intensity. Oocytes were imaged at 10× magnification to visualise GFP expression. Created in https://BioRender.com

While oocytes expressing Kesv 1 showed low levels of autofluorescence, the pronounced fluorescence signal concentrated near the equatorial band in Kesv 1-GFP expressing oocytes, indicating a distinct expression pattern compared with the control. Also, in comparison with Kesv 2-GFP, a sharp green fluorescence with higher intensity was observed near the equatorial band in oocytes expressing Kesv 1-GFP tagged constructs, while oocytes bearing the Kesv 2-GFP tags exhibited weaker fluorescence in comparison (Fig. 4 B, C). Further analysis based on different intensities of fluorescence excitation light revealed that although, the exposure time may have differed depending upon the different relative intensity of excitation light under study, a slight difference was observed in terms of emitted fluorescence intensities where 40% excitation (Fig. 4B) showed slightly better expression. The expression pattern followed the trend Kesv 1-GFP > Kesv 2-GFP > Kesv 1, indicating a higher expression of Kesv 1-GFP compared to Kesv 2-GFP and Kesv 1.

### Kesv 1 and Kesv 2 AlphaFold models are similar

To aid the interpretation of the biochemical experiments, we first predicted the monomeric channels using AlphaFold 3 (AF3). The core transmembrane helices (M1, P and M2), and the SF geometry were similar in both channels, while the N-terminal helix adopted different orientations (Fig. 5 A, B). We next modeled the tetrameric structures of the Kesv 1 and Kesv 2 ion channels and again, N-terminal helices adopted a range of orientations (Fig. 5 C, D). For each channel, AF3 generated a set of five homotetrameric models (ranks 0–4). Per-residue confidence scores mapped onto the AF3 rank 0 models show that the pore region, including helices M1, P and M2 and the SF (SVGYG motif), is predicted with uniformly high confidence, whereas the N-terminal helix has lower confidence (Fig. 5 A-D).

**Figure 5.**
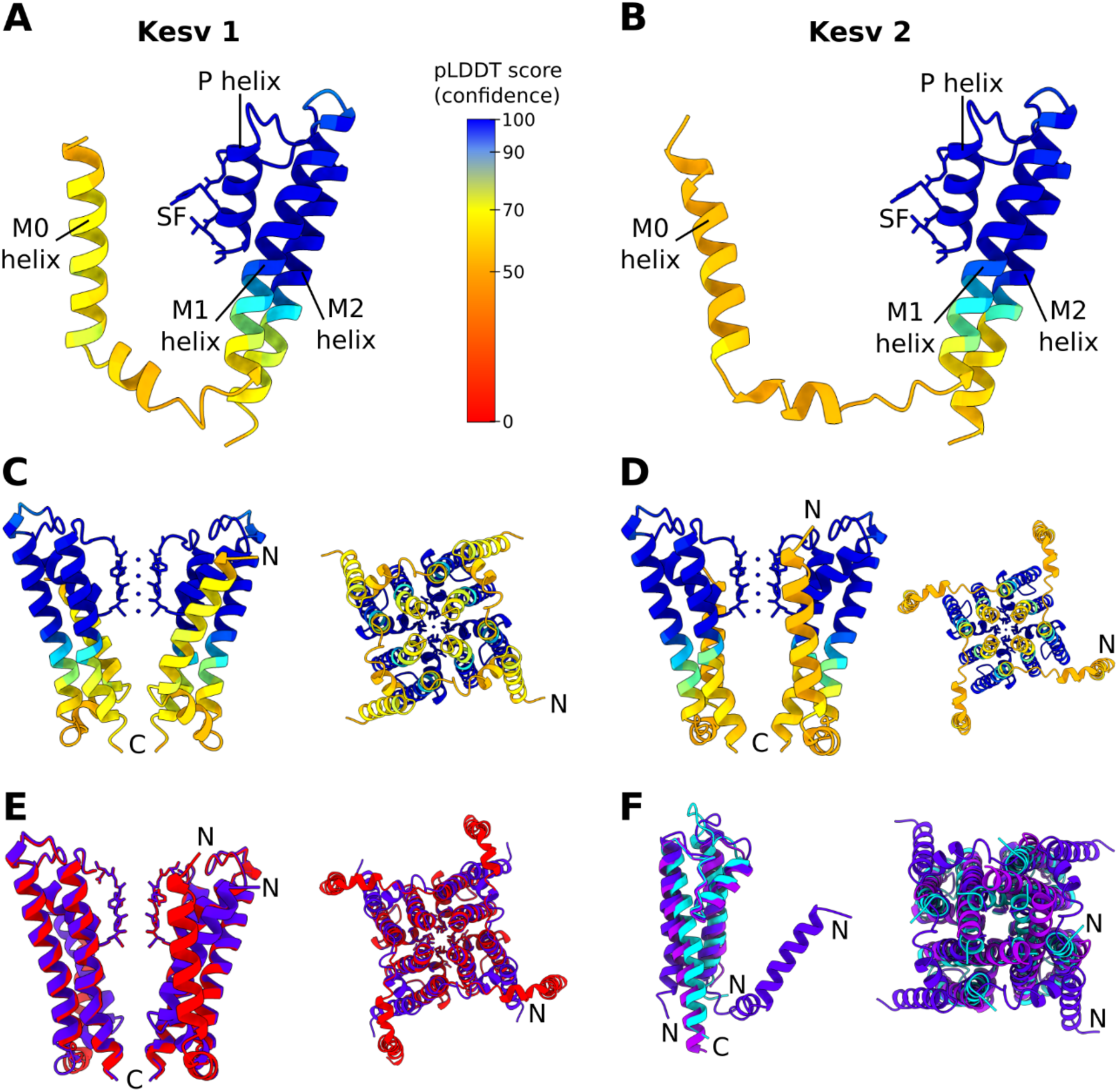
Structural analysis and comparison of Kesv 1 and Kesv 2 using AlphaFold models. (A,B) Monomeric structural predictions of (A) Kesv 1 and (B) Kesv 2 generated with AlphaFold3, coloured by per-residue confidence (pLDDT). (C,D) Tetrameric AlphaFold3 models (model 0) of (C) Kesv 1 and (D) Kesv 2, shown in front (left) and side (right) views and coloured by pLDDT score (low confidence <50; high confidence >90). (E, F) Structural superpositions of Kesv 1 (blue) and Kesv 2 (red) (E), and comparison of Kesv-1 with closed conformations of MthK (PDB 6U5R, purple) and KcsA (PDB 5J9P, cyan) (F) The predicted Kesv structures reproduce the conserved pore-domain architecture and closely match the selectivity filter, with differences mainly in the N-terminal helix.

Structural superposition of Kesv 1 and Kesv 2 (Fig. 5E) shows that the N-terminal ⍺-helix in Kesv 1 bends around residues 20–23, followed by a loop that redirects the helix toward the pore axis. The Kesv 2 models also display a bend in this region, but it is less pronounced and the loop guides the helix along a different path. Thus, AF3 predicts that Kesv 1 and Kesv 2 share a common bent-helix feature but differ in the exact orientation of their N-terminal segments. The five AF3 models of Kesv 1 and Kesv 2 are depicted in Supplementary Figures S1 and S2, PDB files are provided as Supporting Information.

To compare the models quantitatively, we calculated the backbone RMSD of each model, w.r.t. the other four models of the same channel, as well as w.r.t. the models of the other channel (SI Fig. S3). These RMSD calculations were performed for the whole protein as well as only for the core region (i.e. without the N-terminal helices, residues 1-35). The results support the findings of the visual superimposition of the structures. In the core region, all ten AF3 models for Kesv 1 and Kesv 2 are highly similar. At both the whole-protein and core-region levels, the structural differences between each Kesv 1 and all Kesv 2 models, and vice versa, are not substantially higher than the structural variances between the five models of each channel.

To interpret the AF3 models in the context of known K⁺ channels, distances between representative residues on opposite monomers were calculated and compared to average distances in PDB structures of open and closed conformations of the well-characterized K⁺ channels MthK (29 % sequence identity and 48 % similarity to Kesv 1, 28 % identity and 47 % similarity to Kesv 2) and KcsA (22 % identity and 49 % similarity to Kesv 1, 23 % identity and 49 % similarity to Kesv 2) (Fig. 5F, SI Fig. S4, SI Table S11). The distance analysis strongly suggests that the Kesv 1 and Kesv 2 models predicted by AF3 represent closed conformations. A structural superimposition (SI Fig. S4F) reveals that the M1, M2 and P helices and the SF of the Kesv models closely align with closed conformations of MthK and KcsA, with the SVGYG motif in Kesv corresponding to the TVGYG motif in reference structures.

Together with the geometric validation of the AF3 models (SI Table 1), these observations support the use of all five AF3 models as starting points for subsequent atomistic molecular dynamics simulations of membrane-embedded Kesv 1 and Kesv 2.

### Kesv 1 and Kesv 2 models are structurally stable in MD simulations

To assess the structural stability of the AF3 models of Kesv 1 and Kesv 2 in a membrane environment, we embedded the channels in a lipid bilayer and carried out all-atom MD simulations (see Methods). To assess structural integrity of the channels, we monitored the time evolution of the backbone RMSD for all 30 MD trajectories (5 AF3 ranks × 3 simulations per channel, 500 ns each; total simulation time 15 µs). Across all systems, the protein structures remained stable on the simulation timescale (SI Fig. S5, S6). Analyzing the RMSD separately for the Kesv core region and the N-terminal helices indicated that the largest deviations occur in the N-terminal helices, whereas the protein core region remained comparatively stable, in line with the structural variances between the AF3 models (see above).

Regarding ion and water occupancy of the SF, one of the four K^+^ ions initially placed in the SF immediately left the SF in the simulation trajectories. Across all simulation replicas and models, the SF was permanently occupied by at least two K^+^ ions, and in most cases by three. Depending on the number of ions, two to four water molecules additionally occupied the SF. The observed SF occupation and ion movements within the SF are exemplarily illustrated in SI Fig. S7 and consistent with mechanisms discussed by Mironenko *et al.* (27).

Taken together, the MD simulations support the notion that both Kesv 1 and Kesv 2 form tetrameric ion channels that are structurally stable in a membrane environment over microsecond timescales. Observing ion conduction would require a conformational transition from the closed conformation modeled by AF3 to an open (conductive) conformation, which was not observed during our MD simulations.

## Discussion

This study aimed to characterize the two variants of a potassium ion channel protein that despite possessing the same residues within their selectivity filter, exhibited differential ion conductance properties. While Kesv 1 is a potassium ion channel encoded by EsV −1 virus (10), Kesv 2 is the integrated version of Kesv 1 encoding gene into the host algae *Ectocarpus siliculosus* (25). It is believed that, during the lysogeny, EsV-1 completely or partially incorporates into the genome of algae and is maintained in a lysogenic phase for many years (8). Kesv 2 is therefore likely the result of acquired mutations during this process of evolution. Similar mechanisms of integration are also observed in other classes of viruses (28); however, the factors driving the introduction and retention of these gene variants remain a subject of interest. One possible explanation is that certain viruses exist in a co-evolutionary relationship with their host, where viral evolution is shaped by constraints imposed by host cellular environments (29). In such cases, viral protein evolution is strongly influenced by structural and functional requirements, while only a subset of mutations is tolerated (6, 30). Unfortunately, the lack of sufficient sequence data restricts our understanding of evolution within a host population as of now but represents an interesting point to focus on as to why mutations were introduced in the viral sequence integrated into the host algae. It would be interesting to understand whether these gene variants are part of the biological mechanisms in algae and whether they have evolutionary significance.

The analysis of the primary sequence of these channels revealed seven altered residues. Among them, four mutations involved the replacement of one residue to another with similar chemical properties (Alanine to Valine, Valine to Alanine and Leucine to Methionine); two mutations involved the change of a neutral amino acid to a non-polar amino acid (Threonine to Alanine and Threonine to Valine) and one mutation showed replacement of a neutral amino acid with an amino acid with basic properties (Glutamine to Histidine). In general, a single amino acid exchange can influence the gating properties of an ion channel and can shift the I-V relationship (31). Efforts were made to identify such critical mutations to better understand these channels.

While the functional expression of Kesv 1 was already explored by Chen *et al.* (10), this is the first study to characterize the Kesv 2 channel and highlight the differences between Kesv 1 and Kesv 2, on the basis of structural and functional parameters. Significantly higher currents observed for Kesv 1 and abolished currents associated with mutation within the selectivity filter were in accordance with the previous findings (10). Although in some cases, Y89A, albeit not significantly, did show slightly higher currents than uninjected which could be based on batch variability, increased permeability of other ions resulting in higher currents or shift in permeation mechanisms (32). Electrophysiologically, the major functional change was observed with mutations Q61H and T66A in combination. The expression pattern of GFP-tagged channels was also consistent with the electrophysiological findings when higher expression of Kesv 1-GFP plasmid was observed. This indicates that both Kesv 1 and Kesv 2 are functionally expressed in oocytes, although Kesv 1-GFP shows more intense, localized fluorescence compared to Kesv 2-GFP.

The impact of these mutations on different localizations within the cell was also highly speculated based on existing literature suggesting the expression of Kesv on mitochondrial membrane (33) in contrast to plasma membrane expression of similar proteins (34). Therefore, the presence of mitochondrial transit peptide (mTP) or signal peptide (SP) for differential subcellular location (mitochondrial or secretory pathway) were investigated using the TargetP-2.0 tool (SI Fig. S7) (35). A putative N-terminal cleavage site was predicted for both variants between residues 19 and 24 with a higher cleavage-site confidence for Kesv 1. A sharp, narrow peak starting at residue Val20 appears for Kesv 1, whereas Kesv 2 shows a broader peak with a higher local probability at Ala20. Since, Kesv is a small membrane protein whose N-terminus corresponds to the first transmembrane helix, this prediction only reflects the similarity between signal peptides and N-terminal transmembrane helices rather than true proteolytic cleavage. (SI Fig. S8). Further, the probability of mTP was low (0.02 for Kesv 1 and 0.04 for Kesv 2) suggesting no direct evidence of mitochondrial expression of this protein (SI Table S12).

The N-terminal truncation of Kesv 1 resulted in the complete loss of ion channel activity, indicating that the N-terminus is essential for channel function. A similar observation was reported for the viral K^+^ channel Kcv, in which deletion of the N-terminus abolished channel activity without affecting protein synthesis or subcellular localization, suggesting that the N-terminus directly contributes to channel function (36). In contrast, N-terminal truncations of the ion channels NaK and MthK increased channel activity, probably by increasing the probability of open conformations in the absence of the N-terminii (37–39). In MthK, the N-terminus was shown to inactivate the channel through a *ball-and-chain* inactivation mechanism (37). Taken together, these findings demonstrate that the N-termini of ion channels – reported to be the least conserved parts of the sequence across different organisms – can fulfill remarkably diverse functions. Kesv 1 represents another example emphasizing that the roles of K^+^ channel N-termini remain poorly understood and warrant further investigation.

Compounds like Sotalol, Linopirdine, and Retigabine at different μM concentrations were chosen for this study based on their ability to regulate potassium channel function in eukaryotes (40). Retigabine is reportedly an activator of Kv 7 potassium channels that increases ion conductance by shifting the voltage-dependence of channel activation to hyperpolarising potentials (41). The key binding sites of Retigabine were previously identified in Kv 7.3 as Trp-265, Gly-340, and Leu-314 within the pore region. Later, Lange et al., (42) refined the binding sites and mode of action of Retigabine on Kv 7.3 structure by identifying other contributors and interactions, notably of Trp-265 with Leu-314 for mediating the activation effect. In this study, Retigabine showed no effect on ion conductance of Kesv 1 channel, which could be attributed to the following reasons. Firstly, as discussed, Retigabine interacts with specific residues and thus acts precisely on channels with a specific architecture (42). Secondly, differences in intracellular signaling, drug concentrations, and inadequate drug access could also potentially lead to undetectable activity of these compounds. Therefore, the lack of change in conductance observed here highlights the dependence of Retigabine’s efficacy on specificity in channel expression and experimental context.

The investigations with other modulators revealed Sotalol as a weak agonist, while Linopirdine as a strong agonist of the potassium ion channel, based on the magnitude of current enhancement and the concentration required to elicit the pharmacological effect relative to control conditions. Sotalol is a non-selective β-adrenergic blocker that also blocks potassium channels by acting directly on the pore region, preventing the flow of K^+^ ions (43). Linopirdine, on the other hand, is also associated with inhibition of voltage-gated potassium channels (44). Both drugs act by suppressing repolarization, thereby increasing the duration of the action potential.

The observed increased conductance with the application of Sotalol and Linopirdine on Kesv channels might be due to the absence of regulatory domains that change the pore accessibility, while, in human potassium channels, these drug effects are shaped by large regulatory domains that indirectly influence the pore (45). By comparing the drug effects in these two systems, the aspects of drug action derived from the channel itself versus those arising from the regulatory domains can be easily distinguished. Therefore, our findings suggest that the observed effect of activation arises from direct interactions between drugs and the core channel structure and explain how complexity can influence the behavior of channel proteins and the overall mechanisms of pharmacology. Nevertheless, this study underscores the importance of context-and channel-specific pharmacology that could yield future therapeutic insights warranting further studies.

## Conclusion

In order to achieve the functional understanding of the Kesv channel protein, the two orthologous sequences of Kesv (Kesv 1 and Kesv 2) were compared to gain insights into their functional expression and pharmacological behaviour. Overall, Kesv 1 showed outward rectifier currents through an active ion channel when expressed in the membrane of *Xenopus* oocytes. In comparison, Kesv 2 exerted no significant functional expression. Seven mutations in Kesv 1 resulted in reduced activity of the ion channel where specifically mutations in close vicinity to the pore helix (Kesv 1-Q61H & Kesv 1-T66A) significantly contributed to the reduced function. All other amino acid exchanges, like Kesv 1 – V20A, Kesv 1 -T103V, and Kesv – Q61H, were found to be functionally preserved, although the amplitude of currents was lower to some extent. The electrophysiological analysis was corroborated by GFP expression of Kesv 1 - GFP and Kesv 2 - GFP in oocytes membrane. The expression was significantly higher for Kesv 1 - GFP than Kesv 2 - GFP, paralleled by a small autofluorescence signal in oocytes expressing only Kesv 1 without the GFP tag. AlphaFold predicts a closed conformation of the channel, with low confidence of the N-terminal M0 helix. Experimentally, the truncation of this helix (residues 1-35) reduced the current amplitudes to those observed for uninjected oocytes, suggesting the importance of the N-terminus in localization, folding, or integration of the channel into the membrane. Taken together, Kesv 1 exhibits all properties of a functionally expressed voltage-gated potassium ion channel. Based on the predicted AlphaFold models, Kesv 1 and Kesv 2 are structurally similar and should in principle both be capable to conduct potassium ions when adopting open conformations, which are currently unknown. Therefore, the experimentally observed differences between Kesv 1 and Kesv 2 are unlikely to be explained by structural differences (as predicted by AlphaFold) alone, but may instead result from differences in expression levels, folding, membrane insertion, or trafficking that reduce the surface expression of Kesv 2.

The interaction of Kesv channel with known modulators such as Sotalol and Linopirdine demonstrates that regulatory subunits may introduce additional complexity to the intrinsic properties of the core channel changing the influence of drugs. These findings suggest that pharmacological effects may be context-dependent, with drug responses differing between the minimal channel (isolated core channel) and the channel with regulatory subunits. The activity of Retigabine, however, emphasized the importance of critical residues around the SF required for drug action. These results may also serve as a useful framework for investigating the evolutionary pressures shaping the elements of viral structure, function, and host–virus interactions.

## Materials and Methods

### Sequence and alignment

A database search was performed to extract the sequence for Kesv protein. The search was performed on databases of NCBI with the following keywords-Kesv, *Ectocarpus siliculosus* virus, potassium ion channel. The nucleotide sequence of Kesv was retrieved (NC_002687.1) (10) and its corresponding amino acid sequence was obtained (Uniprot ID-Q8QN67). The similarity search was then performed by BLASTp and this led to an identification of a new Kesv sequence (Kesv 2) (uniport ID-D8LP37) sharing similarity of ∼94 % with the first sequence Kesv 1 (Accession number-CBN80308.1) (25).

### Molecular Biology and TEVC measurements

Kesv 1 and Kesv 2 were subcloned into psGEM vector according to standard protocols (46). For *in vitro* transcription, DNA templates were linearized with Nhel-HF for cRNA preparation (mMESSAGE mMACHINE T7 Transcription Kit, Invitrogen, Carlsbad, USA). Defolliculated oocytes (EcoCyte Bioscience, Dortmund, Germany) were injected with 20 ng cRNA for each variant. After injection, oocytes were incubated for 2 days in Barth’s solution (88 NaCl mM, 1 mM KCl, 0.4 mM CaCl_2_, 0.33 mM Ca(NO_3_)_2_, 0.6 mM MgSO_4_, 5 mM TRIS-HCl, 2.4 mM NaHCO_3_, supplemented with 80 mg/L theophylline, 63 mg/L benzylpenicillin, 40 mg/L streptomycin and 100 mg/L gentamycin). TEVC recordings were performed according to standard procedures (24) in ND96 solution containing 96 mM NaCl, 2 mM KCl, 1.8 mM CaCl_2_, 1 mM MgCl_2_, and 5 mM HEPES (adjusted to pH 7.4 with 1M NaOH). The recording pipettes were backfilled with 3M KCl (resistance 0.5-1.5 MΩ). For recording, the oocytes were held at a holding potential of −80 mV for 3 s, followed by a one-second test pulse to potentials ranging from −120 to +40 mV. Currents were measured at the end of one-second pulses and plotted against voltage to obtain the current-voltage (I-V) relationship. Data were expressed as means ± SEM (n = number of oocytes).

### Site-directed mutagenesis

To identify the residues responsible for the reduced functional expression of Kesv, the codons of variant amino acids—selected based on their location—were replaced in the Kesv 1 template using site-directed mutagenesis. The variants such as (Q61H & T66A), V20A, Q61H, T103V and Kesv 1-NΔ35 (1-35 N-terminal residue truncation) were selected and site-directed mutagenesis was performed according to a standard protocol (47). Additionally, mutagenesis involving Y89A was also introduced to silence the selectivity filter. For *in vitro* transcription, linearization of the mutants was performed by *Nsi*I and cRNA was prepared (HiScribe T7 ARCA mRNA kit). Defolliculated oocytes (EcoCyte Bioscience, Dortmund, Germany) were injected with 20 ng cRNA for each variant along with the wild type Kesv 1. After injection, oocytes were incubated for 2 days in Barth’s solution and TEVC recordings were performed using the same protocol as stated above.

### Data analysis and statistics

Significance of mean differences were analyzed by one-way-ANOVA and post hoc mean comparison Tukey test indicated by ns for p > 0.05, * for p < 0.05, ** for p < 0.01 and *** for p < 0.001.

### Fluorescence imaging of channel expression

The EGFP-tagged psGEM plamsids were received from Prof. Guiscard Seebohm and defolliculated oocytes (stage V-VI) were obtained from Ecocyte Bioscience, Dortmund. The cloning of Kesv 1 and Kesv 2 genes in EGFP-psGEM plasmids were performed using standard restriction free cloning procedure and plasmid was isolated using high copy number plasmid purification kit from Macherey-Nagel. The oocytes were injected with 20 ng of EGFP Kesv 1/EGFP Kesv 2 cRNA using Micro 4 Syringe Pump Controller UMC-4 (Word Precision Instruments), and the expression was recorded using a fluorescence microscope (BZ-X1000, Keyence, Germany). The oocytes were oriented under 10x magnification such that the animal pole positioned on the left and vegetal pole on the right, separated by an equatorial band (48).

The excitation light at 40% and 100% intensities was used to monitor and compare the intensities of fluorescence emission. Imaging conducted at 40% excitation was performed with a minimum exposure time of 1/1.5 s while 0.25 s was used for 100% excitation. Imaging proceeded from one edge of the oocyte, using quick full focus, systematically capturing fluorescence across the entire cell. The comparisons were made between Kesv 1-GFP, Kesv 2-GFP and Kesv 1 plasmid without GFP tag, serving as a control.

### Pharmacological modulation of Kesv

Compounds like Retigabine, Sotalol and Linopirdine at different μM concentrations were used to assess their pharmacological behaviour on activity of Kesv 1 expressing oocytes. For Sotalol, 1 mM stock solution in ND96 buffer while 10 mM stock solution for Retigabine and Linopirdine were prepared in 99.8% DMSO. Subsequently, working stocks of 5 μM, 10 μM, 20 μM, and 50 μM were prepared for these compounds in 20 ml of ND96 buffer. The two additional concentrations, i.e. 0.1 μM and 1 μM were also used for Linopirdine. The final concentration of DMSO for the working stocks was adjusted corresponding to the highest observed concentration, i.e. 0.5% for 50 μM compound.

Once these stocks were prepared, the injection was performed following the standard protocol (49). For each concentration, 20 oocytes with cRNA of Kesv 1 wild type was injected and incubated in Barth solution for 2 days at 16°C. The effects of these compounds on the activity of Kesv 1 were then analysed using standard electrophysiological TEVC method (26). The control Kesv 1 - injected and uninjected oocytes were also recorded in the presence of ND96 buffer containing 0.5% DMSO. In addition, uninjected oocytes were also exposed to the highest compound concentration used in this study (i.e., 50 µM Sotalol or Retigabine) to rule out the possibility of compound- or solvent-induced effects on endogenous oocyte currents. The I-V data obtained from different batches were normalised to I_max_ on a 0-1 scale. Further statistical tests were performed comparing Kesv 1-expressing oocytes recorded in 0.5% DMSO with Kesv 1 in the presence of each compound at its respective concentration. Differences between Kesv 1-injected and uninjected oocytes in 0.5% DMSO were also statistically analysed.

### Structure prediction

Tetrameric models of the Kesv 1 and Kesv 2 channels were predicted using AlphaFold 3 (AF3) (50). For each channel, the full-length monomer sequence (124 amino acids) was submitted as a homotetrameric complex. AF3 produced five ranked models per channel. For illustration, AF3 per-residue confidence scores were mapped onto the models and visualized as color-coded structures (relative model confidence) (SI Fig. S1 and S2).

The geometric quality of all AF3 models was assessed using the MolProbity web server (51). (SI Table S1). All five ranked AF3 models for each channel exhibited acceptable overall stereochemistry and were therefore retained as starting structures for the subsequent molecular dynamics stability simulations in the next section. (50, 52–54).

### Molecular dynamics simulations

All simulations were performed on the tetrameric Kesv 1 and Kesv 2 models predicted by AF3 (section: Structure prediction). For each channel, all five AF3-ranked tetramers were used as starting structures. In every case, four K⁺ ions were initially placed in the SF.

Initial protonation states were assigned at pH 7 using the APBS software suite (55), and the channels were embedded in a mixed lipid bilayer containing 200 lipids (SI Table S2). System construction (membrane insertion, solvation and ion placement) was carried out with the CHARMM-GUI webserver (56, 57), yielding an overall neutral periodic cubic simulation box with approximately 9.4 nm side length containing TIP3P (58) water molecules as well as K⁺ and Cl⁻ ions at a concentration of 0.15 M.

GROMACS input structures and topology files were generated with CHARMM-GUI, and all simulations were run with GROMACS 2021.5 (59) using the CHARMM36m force field (57) and the CHARMM-specific TIP3P water model (58). After energy minimization with the steepest-descent algorithm, each system was equilibrated in two stages with all protein heavy atoms position-restrained (59). First, a 20 ps NVT equilibration at 300 K was performed, followed by a 100 ns NPT equilibration at 300 K and 1 bar. The velocity-rescale (v-rescale) thermostat (60) was used to maintain constant temperature, and the stochastic cell-rescale (c-rescale) (61) barostat was used in semi-isotropic mode for constant pressure.

Production simulations were performed after removing all position restraints, using the leap-frog integrator with a 2 fs time step. All bonds involving hydrogen atoms were constrained using the linear constraints solver for molecular simulations (LINCS) (62). Van der Waals energies and forces were smoothly shifted to zero between 1.0 and 1.2 nm, and long-range electrostatics were treated with the particle-mesh Ewald (PME) method (63) with a 1.2 nm real-space cutoff and 0.12 nm grid spacing.

For both Kesv 1 and Kesv 2, each of the five AF3 models was simulated in three independent replicas starting from different random seeds for generating the initial atomic velocities at 300 K. Each replica was run for 500 ns, resulting in a total of 30 stability simulations and an aggregate simulation time of 15 μs.

Trajectory analysis was carried out with standard GROMACS tools and Python scripts. RMSD profiles were computed with gmx rms using the equilibrated structure as reference. Secondary-structure time courses were computed with gmx dssp (DSSP) (64) and summarized as residue-wise secondary-structure profiles over time.

## Acknowledgments

This work was supported by Deutsche Forschungsgemeinschaft (DFG) under Germany’s Excellence Strategy - EXC 2033 - 390677874 - RESOLV. PA would like to acknowledge the financial assistance received from German Academic Exchange Service (DAAD), Germany under the programme: Research grants-Doctoral programmes in Germany, 2021/2022.

## Author Contributions

P.A., L.V.S., G.S., R.S. designed research; P.A., A.E., A.P.Z. performed research, P.A., A.E., A.P.Z., L.V.S., G.G., J.A.S., R.B., D.Todt. analyzed data; D. Tapken contributed new reagents/ analytic tools and P.A., R.S. wrote the paper. All authors edited the draft of the manuscript.

## Supporting Information

### Supporting Information Text

**Fig. S1.**
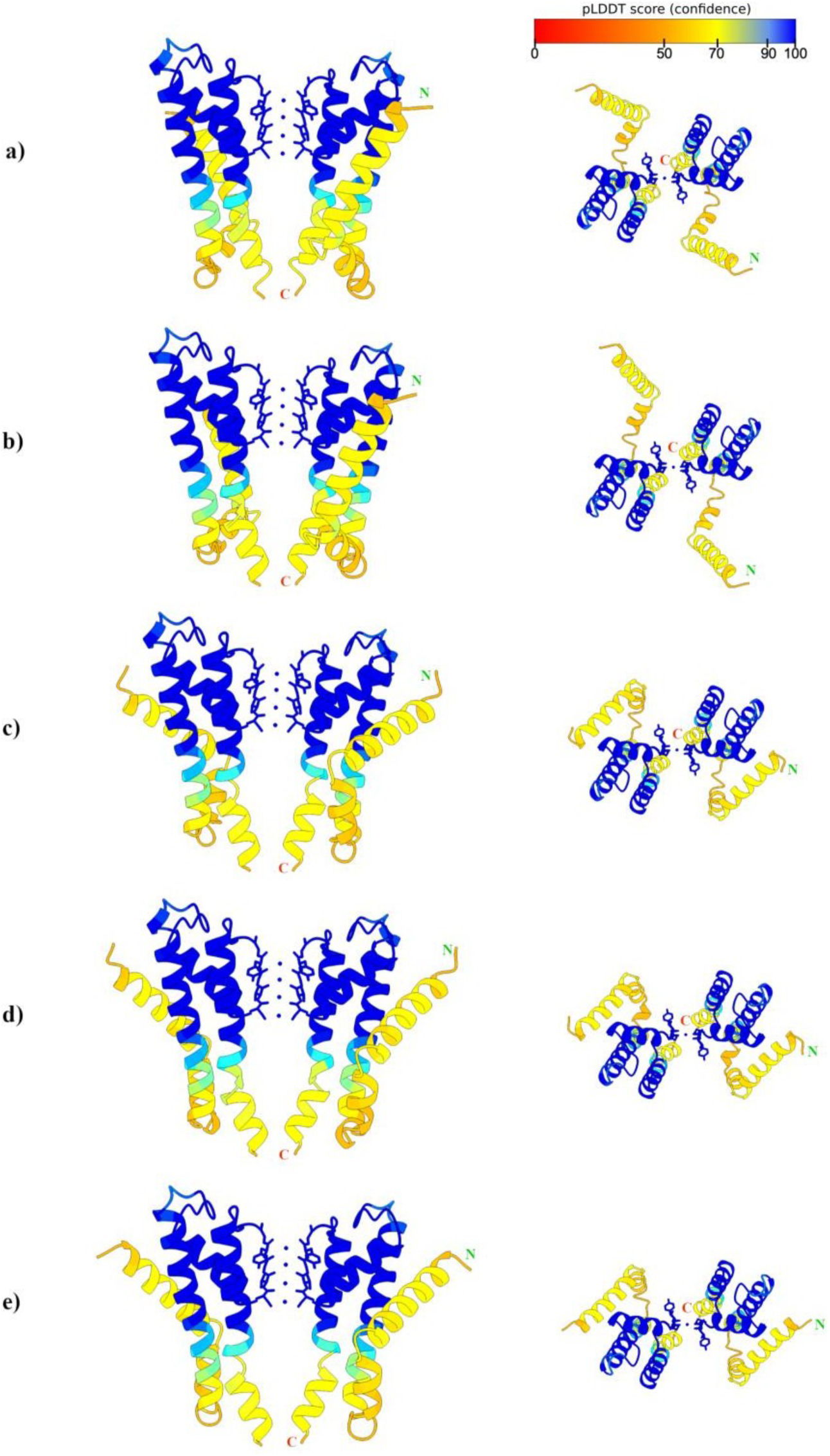
Five AlphaFold3 (AF3) structural models of the Kesv 1 tetramer are shown (a–e, corresponding to models 0–4, respectively). For each model, two views are provided: a front view (left) oriented to visualize the selectivity filter (SF) and pore region, and a top view (right) highlighting conformational differences in the N-terminal helices. Structures are colored by AF3 predicted local confidence, ranging from blue (highest confidence) to orange (lowest confidence). For visual clarity, two subunits were removed from each tetrameric model prior to rendering. Across the model ensemble, the dominant structural variability is localized to the N-terminal helices and the subsequent connecting loop, whereas the overall channel architecture and SF region remain qualitatively conserved.

**Fig. S2.**
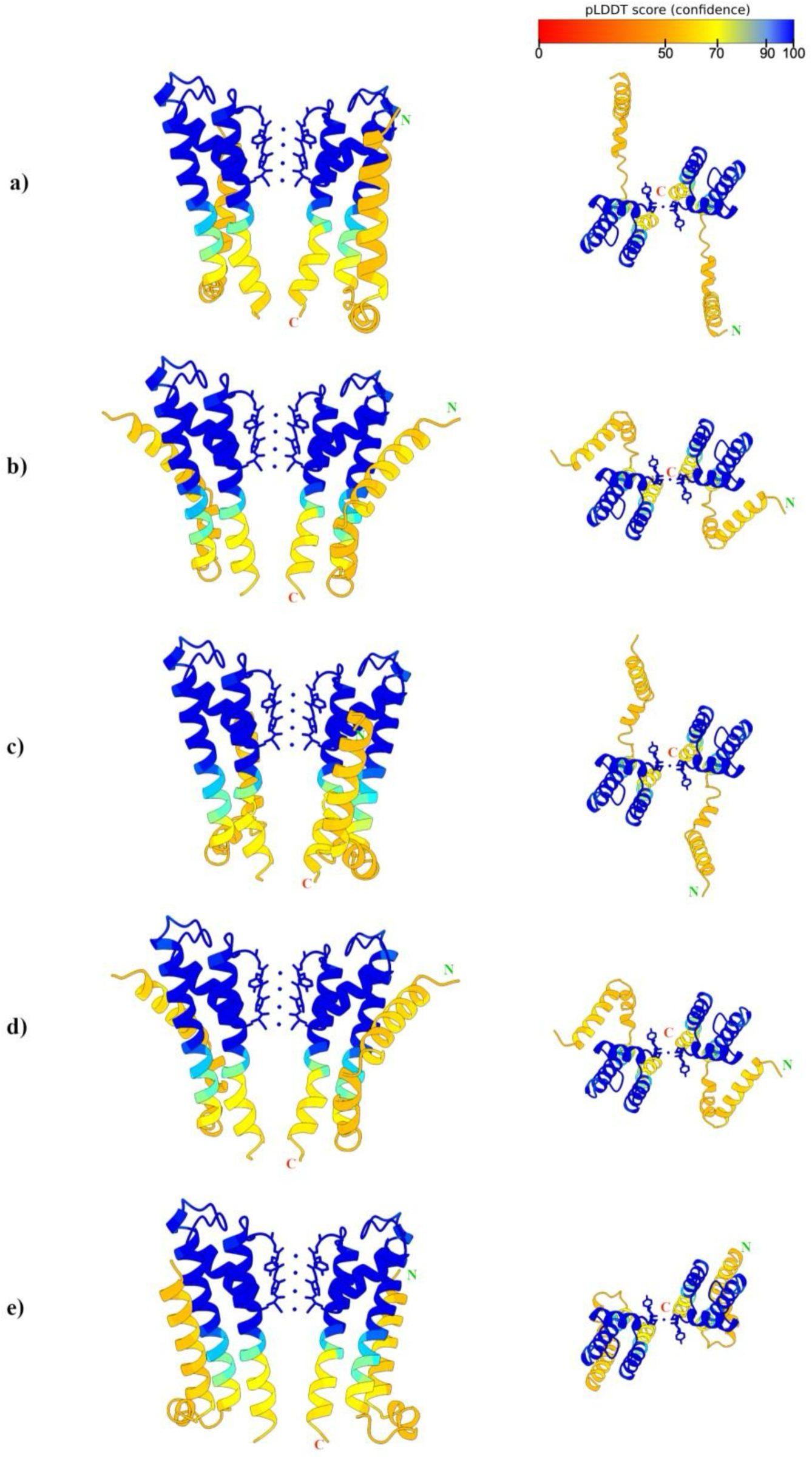
Five AlphaFold3 (AF3) structural models of the Kesv 2 tetramer are shown (a–e, corresponding to models 0–4, respectively). For each model, two views are provided: a front view (left) oriented to visualize the selectivity filter (SF) and pore region, and a top view (right) highlighting conformational differences in the N-terminal helices. Structures are colored by AF3 predicted local confidence, ranging from blue (highest confidence) to orange (lowest confidence). For visual clarity, two subunits were removed from each tetrameric model prior to rendering. Across the model ensemble, the dominant structural variability is localized to the N-terminal helices and the subsequent connecting loop, whereas the overall channel architecture and SF region remain qualitatively conserved.

**Fig. S3.**
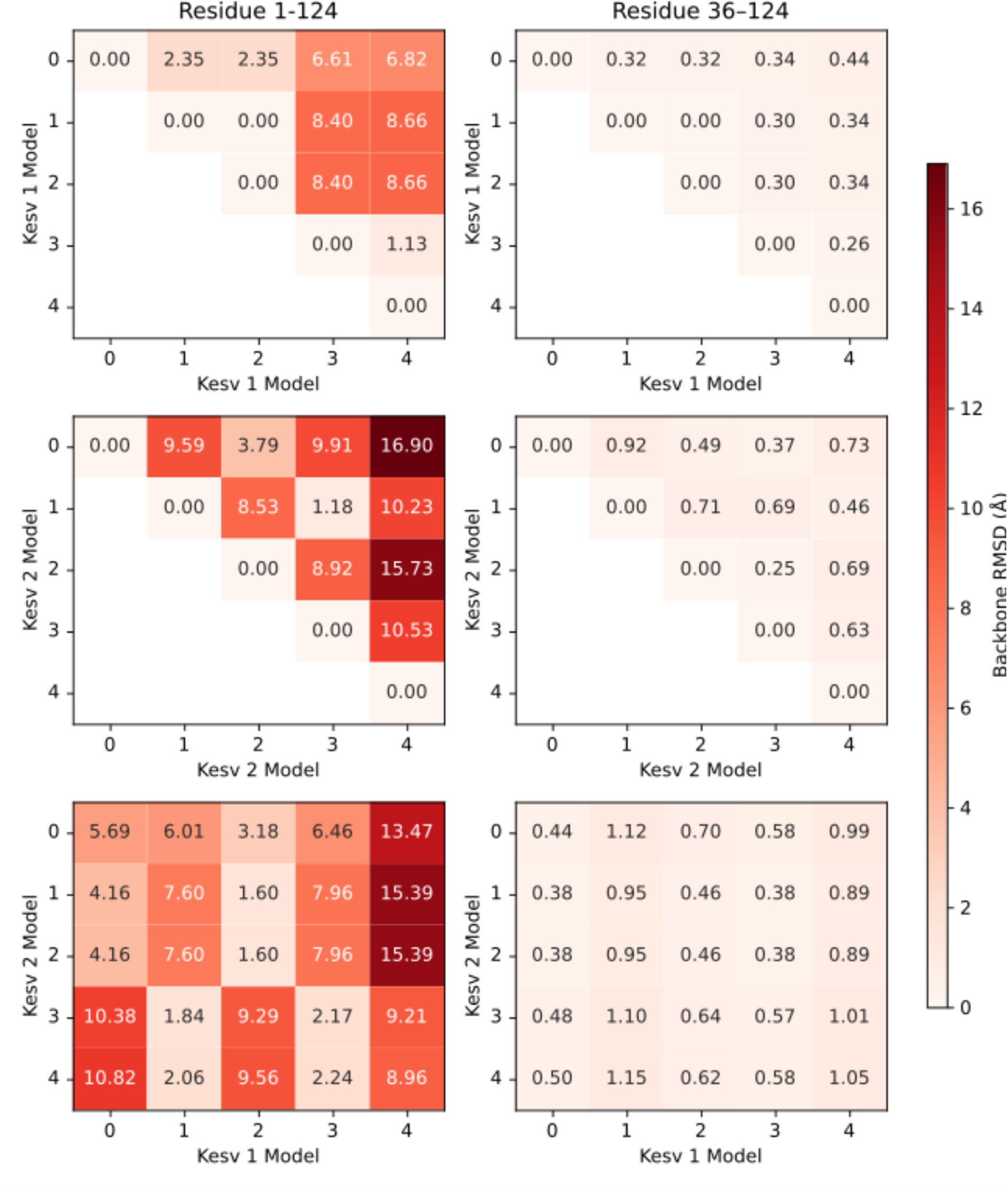
Comparison of Kesv 1 and Kesv 2 models predicted by AF3. The heatmaps indicate the backbone RMSD of each Kesv 1 model w.r.t. the other Kesv 1 models (upper row), of each Kesv 2 model w.r.t. the other Kesv 2 models (middle row) and of each Kesv 1 model w.r.t. each Kesv 2 model. For the data in the left column, the whole tetrameric assembly was considered for analysis, while for the RMSD calculations shown in the right column, the N-terminal helices (residues 1-35) were excluded.

**Fig. S4.**
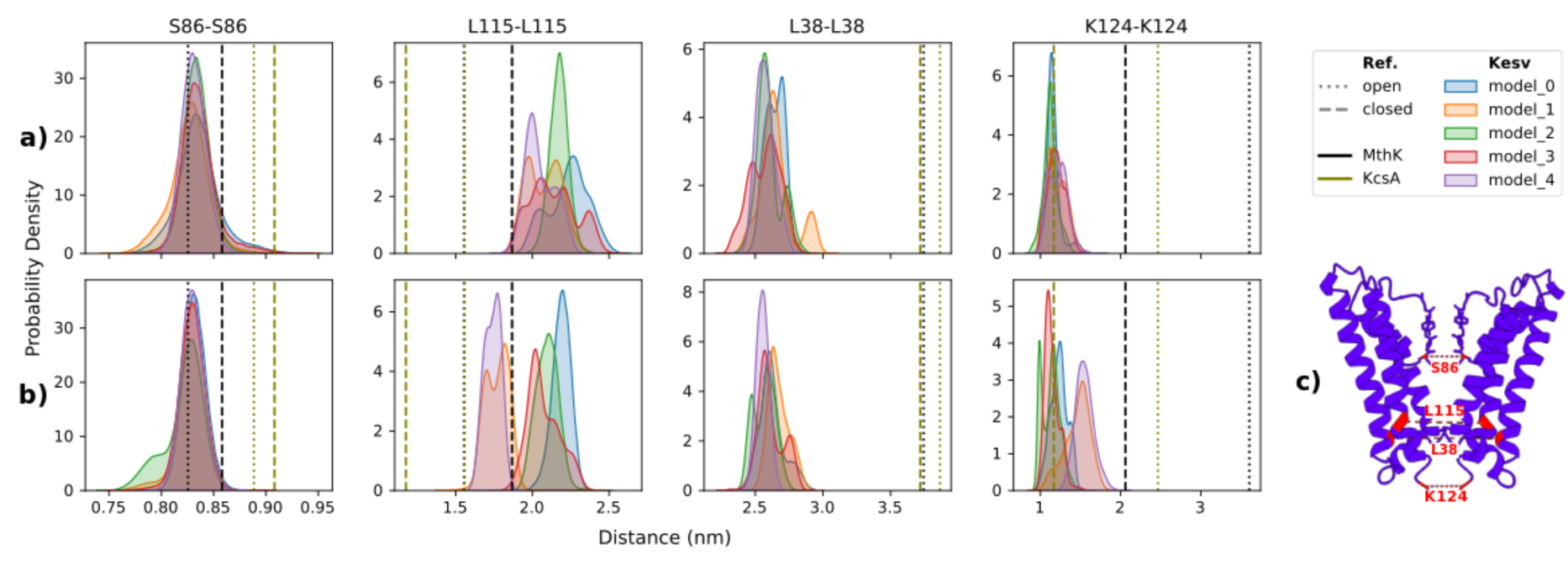
Distribution of distances between C*_α_* atoms of representative residues on opposite monomers throughout the MD simulations of Kesv 1 (a) and Kesv 2 (b) models in comparison to average distances in PDB structures of open and closed conformations of MthK (PDB IDs 6U5R, 8DJB, 5BKI for closed and 1LNQ, 3LDC, 4QE9, 6OLY, 6U9P for open conformations) and KcsA (PDB IDs 1BL8, 1F6G, 5J9P, 2QTO, 3EFF, 7MHR for closed and 3F5W, 5VKE, 7M2J, 7MUB for open conformations). The selected residues S86 (SF), L115 (M2 helix), L38 (M1 helix) and K124 (M2 helix), depicted in (c), correspond to T59, A88, P19, F97 in MthK and T75, G104, A31, T112 in KcsA.

**Fig. S5.**
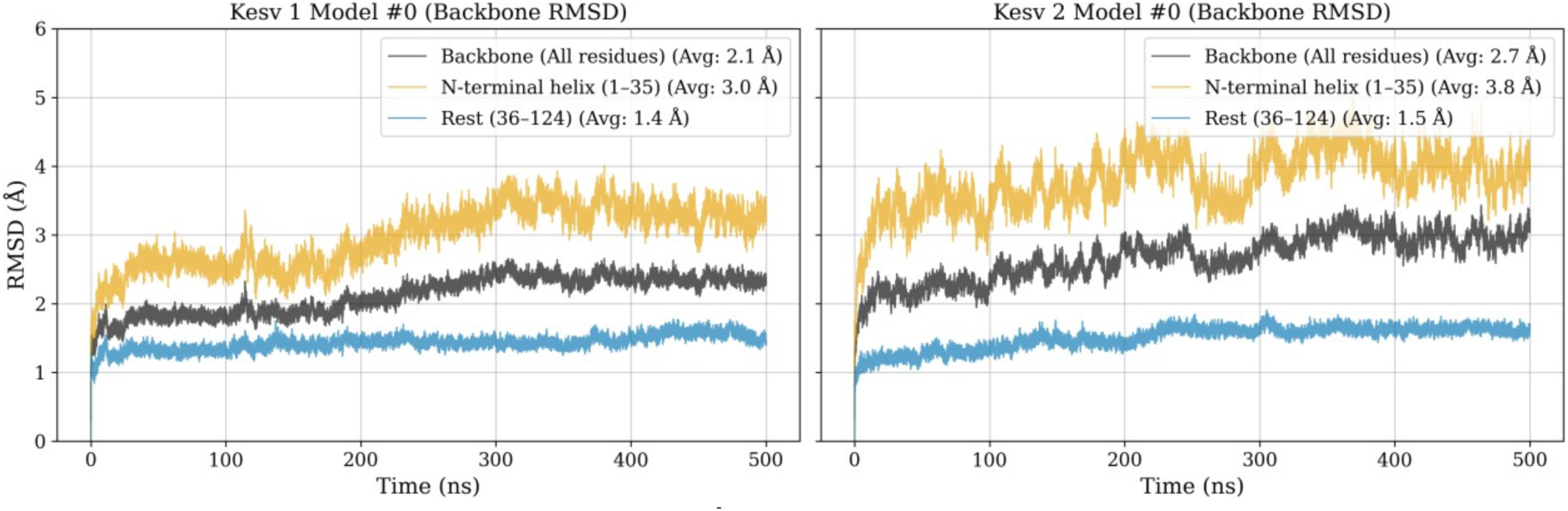
Backbone RMSD time series (Å) are shown for Kesv 1 model #0 (left) and Kesv 2 model #0 (right), calculated with respect to the structure after equilibration. For each channel, RMSD is reported for the full backbone, the backbone of the N-terminal helix (residues 1–35), and the remaining backbone (residues 36–124). In both trajectories, the largest deviations are localized to residues 1–35, whereas residues 36–124 remain comparatively stable, indicating that the overall backbone RMSD is dominated by motion of the N-terminal helix. Mean RMSD values for each trace are reported in the legend.

**Fig. S6.**
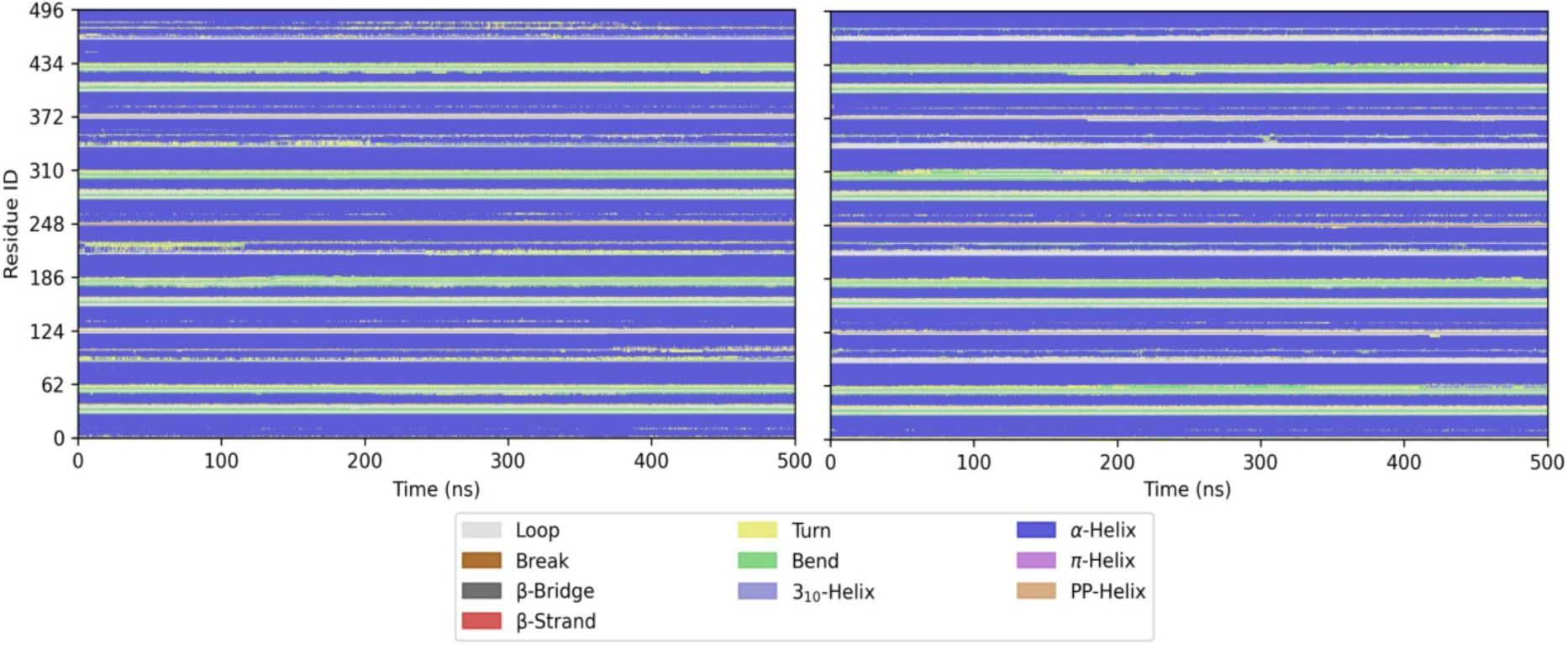
DSSP analysis for Kesv 1 (left) and Kesv 2 (right), indicating the preservation of secondary structure. No significant loss of secondary structure was observed in any of the simulations.

**Fig. S7.**
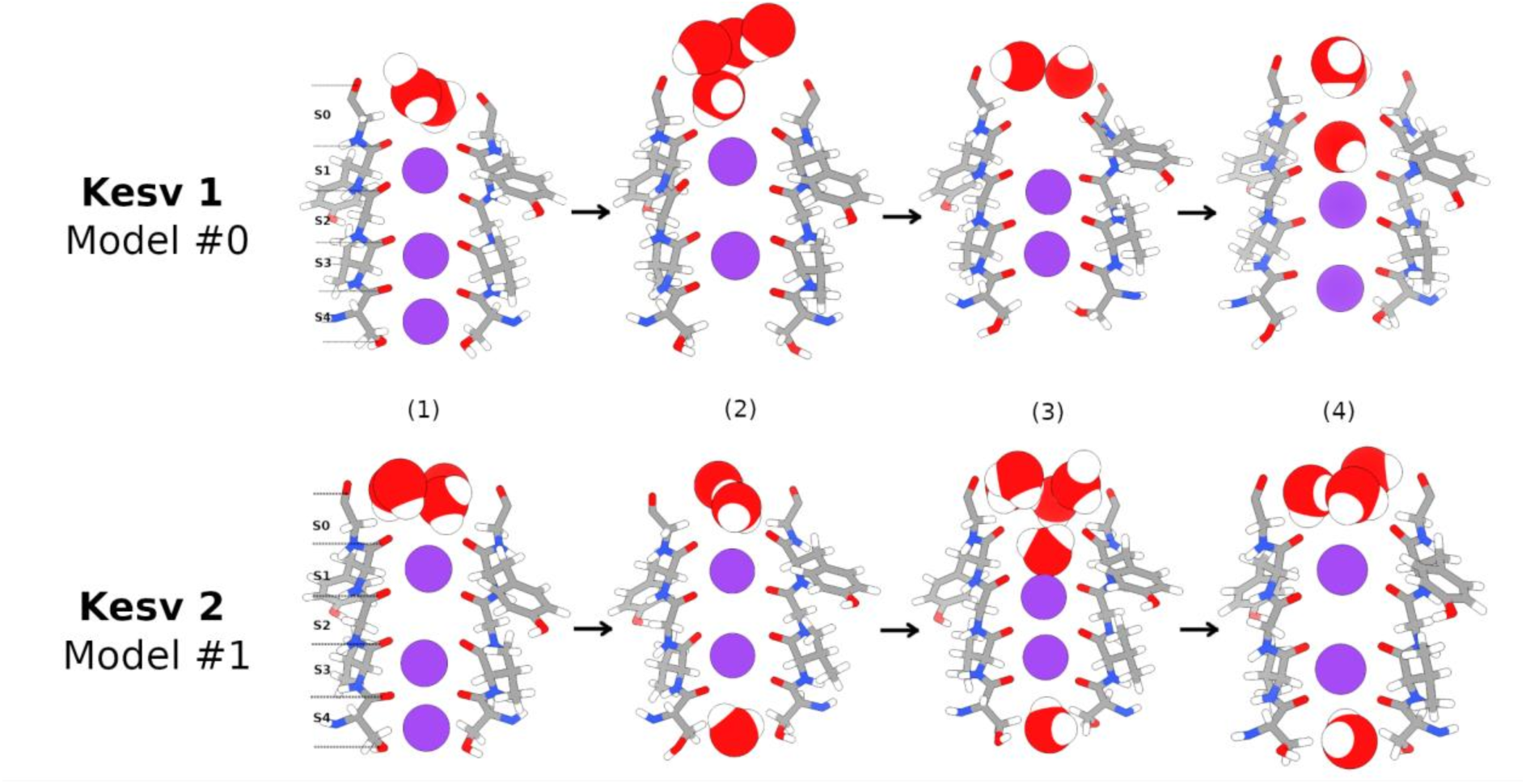
Representative snapshots of the SF illustrating ion movement. The upper panel shows Kesv 1, whereas the lower panel depicts Kesv 2. Columns (1) to (4) represent snapshots of noticeable rearrangements during each simulation. Occupancy at the (S1, S3) and (S2, S4) binding sites persist throughout the simulation. Panel (3) shows an (S3, S2) occupation, which is unfavorable due to electrostatic repulsion and was only observed briefly.

**Fig. S8.**
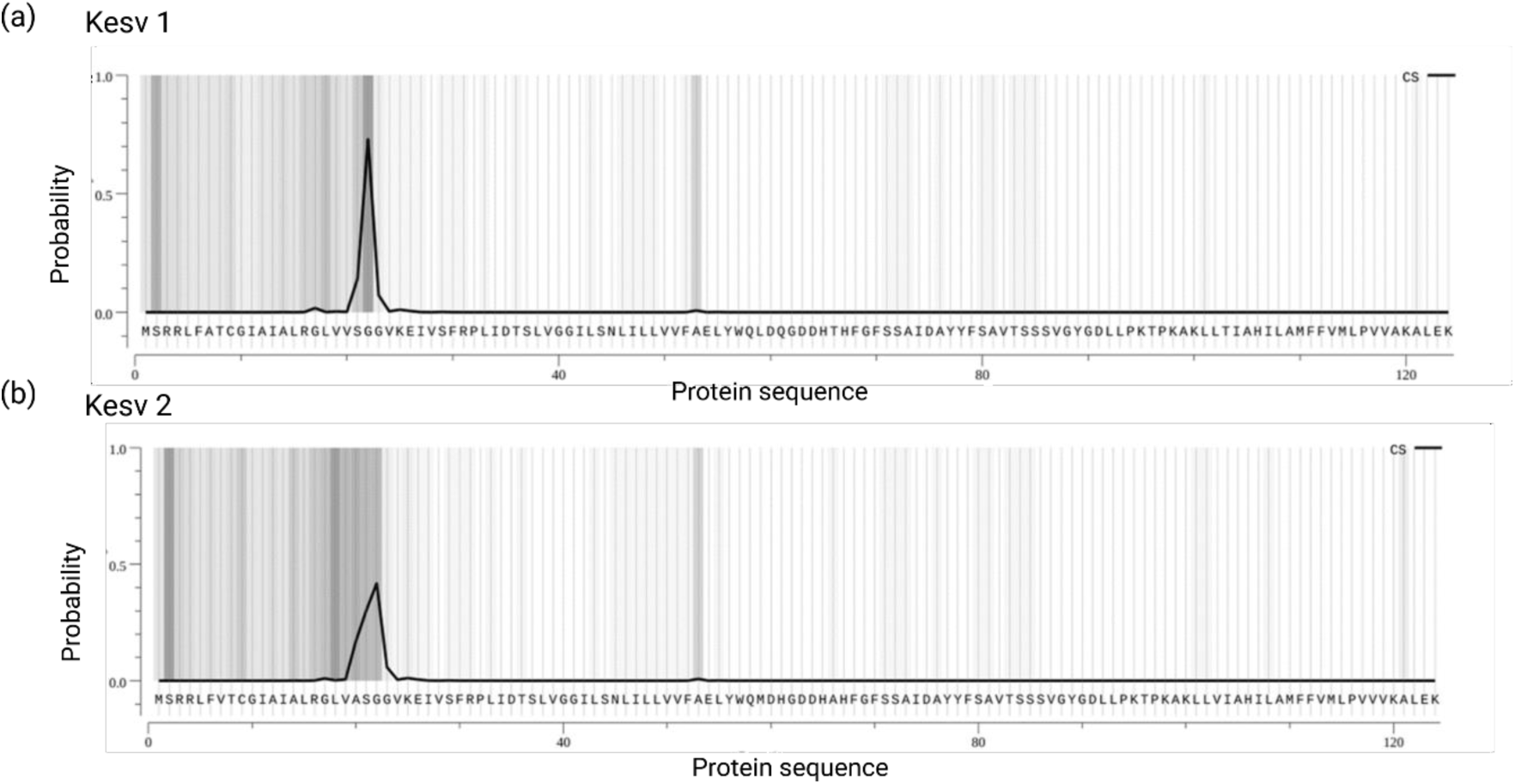
Prediction of plasma membrane targeting sequence of Kesv 1 and Kesv 2 by Target P-2.0 tool.

### Tables

**Table S1.**
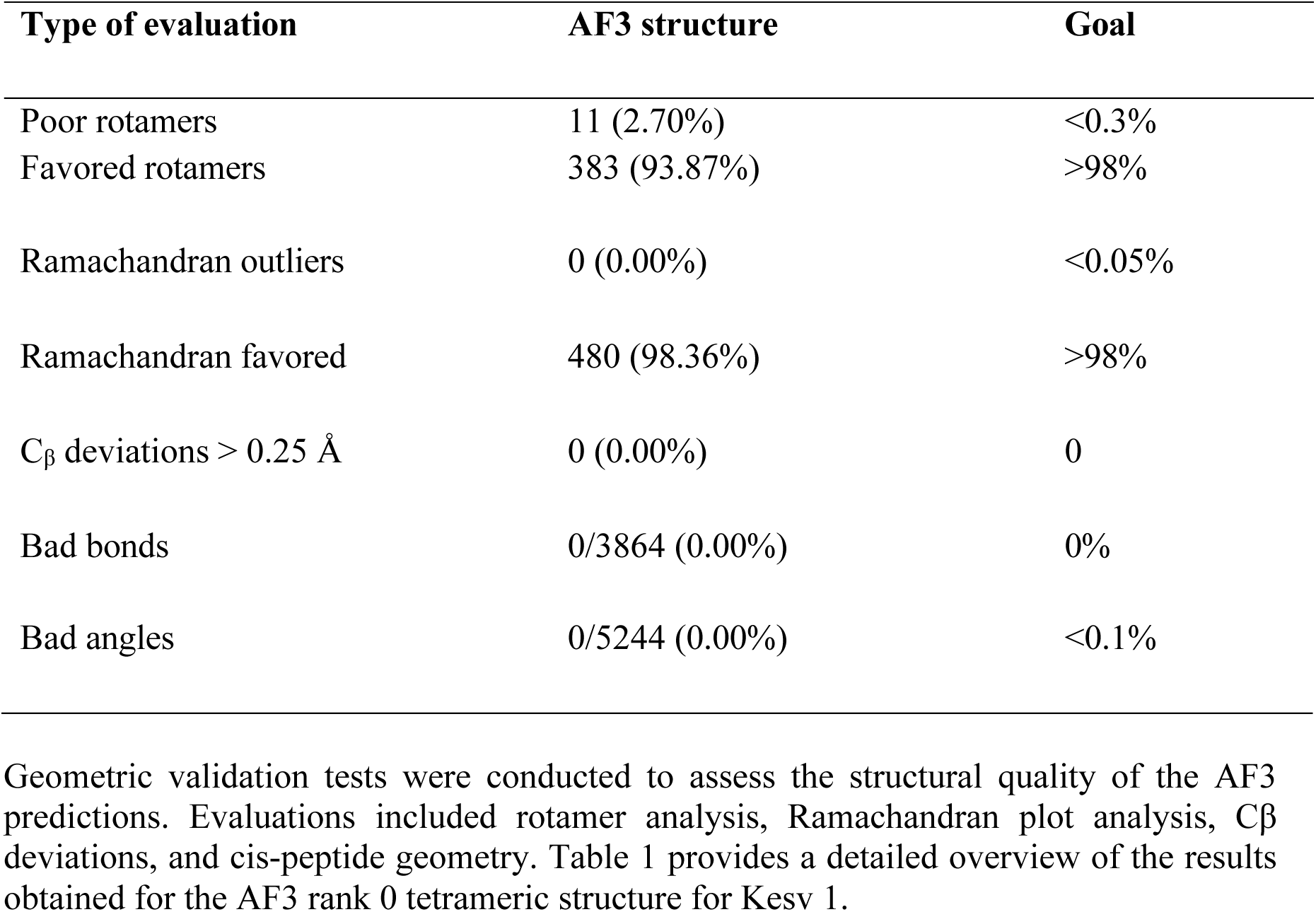
Overview of structural evaluation metrics for the AF3 rank 0 model of Kesv 1.

| Type of evaluation | AF3 structure | Goal |
| --- | --- | --- |
| Poor rotamers | 11 (2.70%) | <0.3% |
| Favored rotamers | 383 (93.87%) | >98% |
| Ramachandran outliers | 0 (0.00%) | <0.05% |
| Ramachandran favored | 480 (98.36%) | >98% |
| Cβ deviations > 0.25 Å | 0 (0.00%) | 0 |
| Bad bonds | 0/3864 (0.00%) | 0% |
| Bad angles | 0/5244 (0.00%) | <0.1% |

**Table S2.**
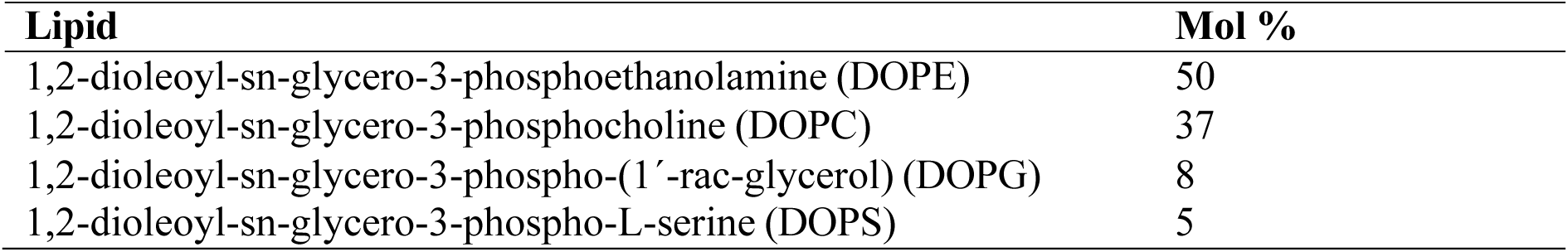
Membrane composition for simulations.

**Table S3.**
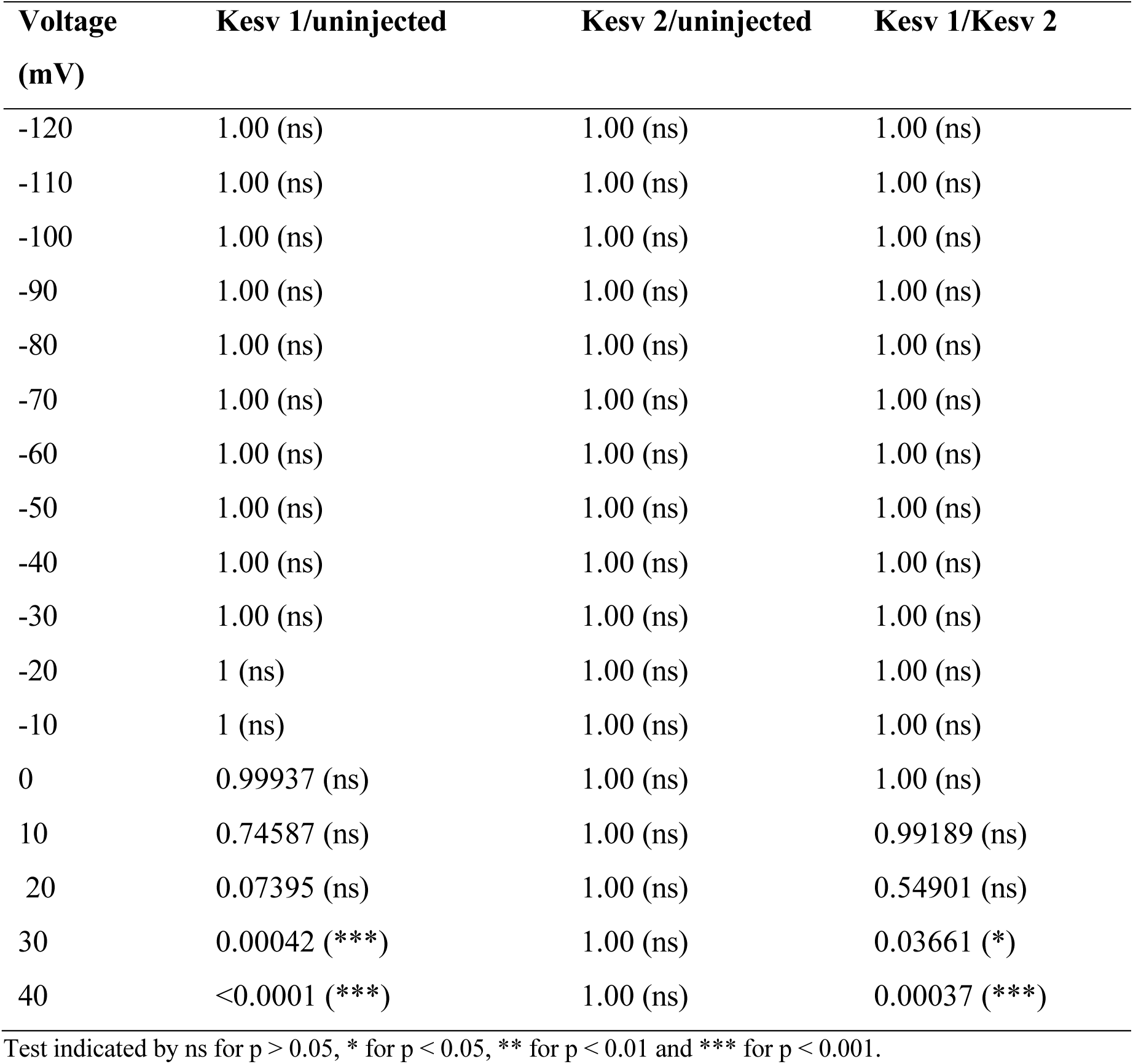
**(a).** p-values for one-way ANOVA with posthoc mean comparison Tukey Test representing functional expression of Kesv 1 and Kesv 2.

**Table S3.** **(b).** Resting potential of *Xenopus laevis* oocytes expressing Kesv 1 and Kesv 2.

| **Construct** | **Number of oocytes** | **Mean resting potential**<br>**[mV ± SEM]** | **P value for comparison**<br>**with uninjected** |
| --- | --- | --- | --- |
| **Kesv 1** | 4 | -55 ± 3.10 | 0.01092 (*) |
| **Kesv 2** | 5 | -49.8 ± 3.15 | 0.05796 (ns) |
| **uninjected** | 4 | -38.5 ± 2.72 | - |

Test indicated by ns for $p > 0.05$ , \* for $p < 0.05$ , \*\* for $p < 0.01$ and \*\*\* for $p < 0.001$ .

**Table S4.**
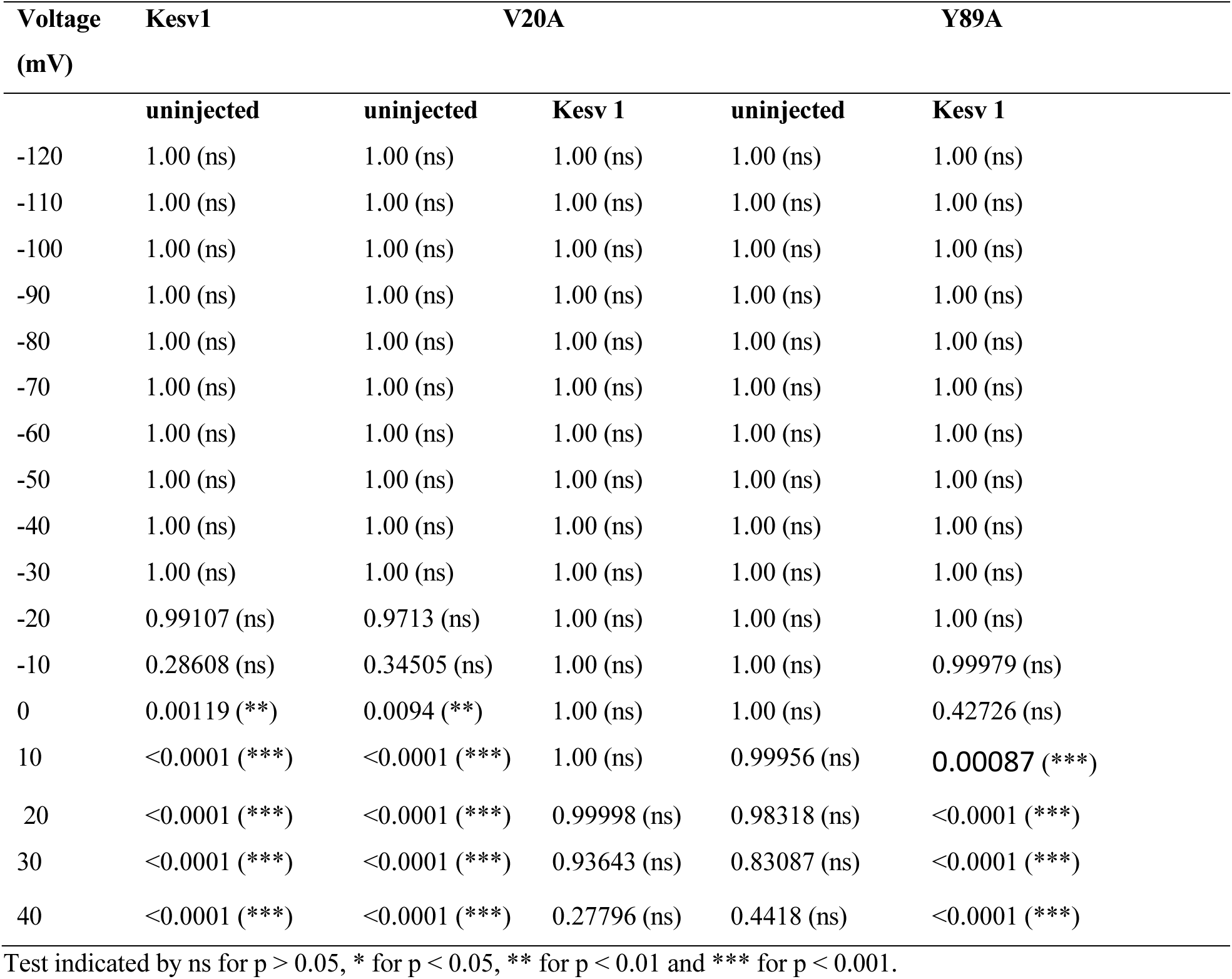
**(a).** p-values for one-way ANOVA with posthoc mean comparison Tukey Test representing functional expression of Kesv 1 (wild type), V20A (mutant), Y89A (negative control).

**Table S4.** **(b).** Resting potential of *Xenopus laevis* oocytes expressing Kesv 1, V20A (mutant), and Y89A (negative control).

| Construct | Number of oocytes | Mean resting potential [mV ± SEM] | P value for comparison with uninjected | P value for comparison with Kescv 1 |
| --- | --- | --- | --- | --- |
| Kescv 1 | 12 | -50.5 ± 4.53 | 0.00156 (**) | - |
| V20A | 12 | -34.58 ± 2.55 | 0.92 (ns) | 0.00605 (**) |
| Y89A | 12 | -27.17 ± 1.79 | 0.96036 (ns) | <0.0001 (***) |
| uninjected | 8 | -30.5 ± 3.74 | - | 0.00156 (**) |

**Table S5.** **(a).** p-values for one-way ANOVA with posthoc mean comparison Tukey Test representing functional expression of Kesv 1 (wild type), Q61H & T66A (mutant), Y89A (negative control).

| Voltage<br>(mV) | Kescv1 | Q61H & T66A |  | Y89A |  |
| --- | --- | --- | --- | --- | --- |
|  | uninjected | Uninjected | Kescv 1 | Uninjected | Kescv 1 |
| -120 | 1.00 (ns) | 1.00 (ns) | 1.00 (ns) | 1.00 (ns) | 1.00 (ns) |
| -110 | 1.00 (ns) | 1.00 (ns) | 1.00 (ns) | 1.00 (ns) | 1.00 (ns) |
| -100 | 1.00 (ns) | 1.00 (ns) | 1.00 (ns) | 1.00 (ns) | 1.00 (ns) |
| -90 | 1.00 (ns) | 1.00 (ns) | 1.00 (ns) | 1.00 (ns) | 1.00 (ns) |
| -80 | 1.00 (ns) | 1.00 (ns) | 1.00 (ns) | 1.00 (ns) | 1.00 (ns) |
| -70 | 1.00 (ns) | 1.00 (ns) | 1.00 (ns) | 1.00 (ns) | 1.00 (ns) |
| -60 | 1.00 (ns) | 1.00 (ns) | 1.00 (ns) | 1.00 (ns) | 1.00 (ns) |
| -50 | 1.00 (ns) | 1.00 (ns) | 1.00 (ns) | 1.00 (ns) | 1.00 (ns) |
| -40 | 1.00 (ns) | 1.00 (ns) | 1.00 (ns) | 1.00 (ns) | 1.00 (ns) |
| -30 | 1.00 (ns) | 1.00 (ns) | 1.00 (ns) | 1.00 (ns) | 1.00 (ns) |
| -20 | 0.99107 (ns) | 1.00 (ns) | 1.00 (ns) | 1.00 (ns) | 1.00 (ns) |
| -10 | 0.28608 (ns) | 1.00 (ns) | 0.99991 (ns) | 1.00 (ns) | 0.99979 (ns) |
| 0 | 0.00119 (**) | 0.99997 (ns) | 0.77434 (ns) | 1.00 (ns) | 0.42726 (ns) |
| 10 | <0.0001 (***) | 0.97274 (ns) | 0.03105 (*) | 0.99956<br>(ns) | 0.00087<br>(***) |
| 20 | <0.0001 (***) | 0.50066 (ns) | <0.0001 (***) | 0.98318<br>(ns) | <0.0001 (***) |
| 30 | <0.0001 (***) | 0.05153 (ns) | <0.0001 (***) | 0.83087<br>(ns) | <0.0001 (***) |
| 40 | <0.0001 (***) | 0.00092 (***) | <0.0001 (***) | 0.4418 (ns) | <0.0001 (***) |

Test indicated by ns for $p > 0.05$ , \* for $p < 0.05$ , \*\* for $p < 0.01$ and \*\*\* for $p < 0.001$ .

**Table S5.** **(b).** Resting potential of *Xenopus laevis* oocytes expressing Kesv 1, Q61H & T66A (mutant), Y89A (negative control).

| Construct | Number<br>of<br>oocytes | Mean resting<br>potential<br>[mV ± SEM] | P value for<br>comparison with<br>uninjected | P value for comparison with<br>Kescv 1 |
| --- | --- | --- | --- | --- |
| Kescv 1 | 12 | -50.5 ± 4.53 | 0.00156 (**) | - |
| Q61H & T66A | 10 | -34.4 ± 3.20 | 0.93996 (ns) | 0.00876 (**) |
| Y89A | 12 | -27.17 ± 1.79 | 0.96036 (ns) | <0.0001 (***) |
| uninjected | 8 | -30.5 ± 3.74 | - | 0.00156 (**) |

**Table S6.**
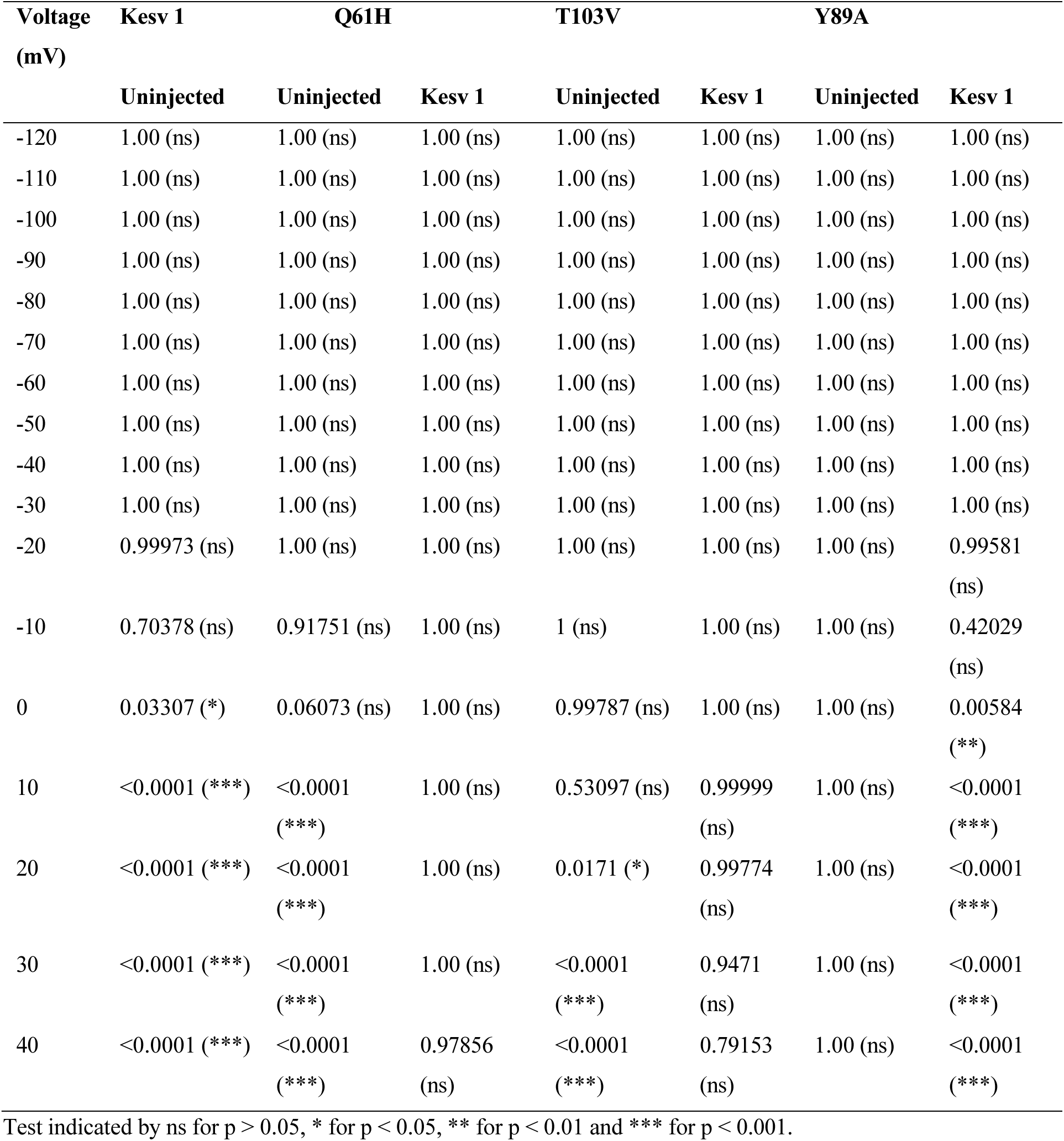
**(a).** p-values for one-way ANOVA with posthoc mean comparison Tukey Test representing functional expression of Kesv 1 (wild type), mutants (Q61H, T103V) and Y89A (negative control).

**Table S6.** **(b).** Resting potential of *Xenopus laevis* oocytes expressing Kesv 1, (wild type), mutants (Q61H, T103V) and Y89A (negative control).

| Construct | Number of oocytes | Mean resting potential [mV ± SEM] | P value for comparison with uninjected | P value for comparison with Kev 1 |
| --- | --- | --- | --- | --- |
| **Kev 1** | 15 | -33.47± 1.47 | <0.0001 (***) | - |
| **Q61H** | 10 | -39.8 ± 1.38 | <0.0001 (***) | 0.2137 (ns) |
| **T103V** | 7 | -32 ± 3.0 | <0.0001 (***) | 0.99878 (ns) |
| **Y89A** | 5 | -6.2 ± 0.97 | 1.00 (ns) | <0.0001 (***) |
| **uninjected** | 4 | -6.75± 2.25 | - | <0.0001 (***) |

Test indicated by ns for $p > 0.05$ , \* for $p < 0.05$ , \*\* for $p < 0.01$ and \*\*\* for $p < 0.001$ .

**Table S7.**
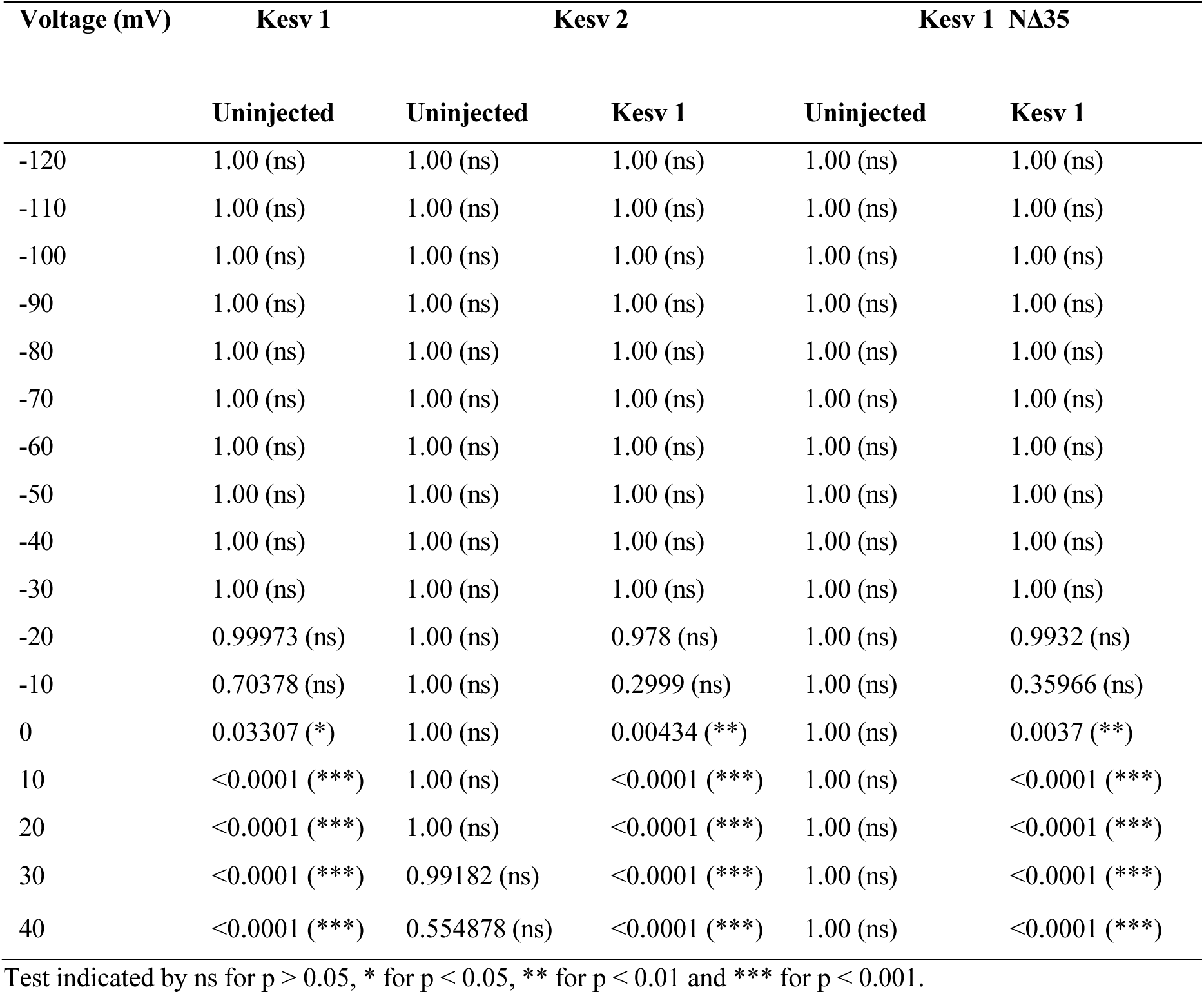
**(a).** p-values for one-way ANOVA with posthoc mean comparison Tukey Test representing functional expression of Kesv 1 (wild type), Kesv 2 and Kesv 1-NΔ35.

**Table S7.** **(b):** Resting potential of *Xenopus laevis* oocytes expressing Kesv 1 (wild type), Kesv 2 and Kesv 1-NΔ35.

| Construct | Number of oocytes | Mean resting potential [mV ± SEM] | P value for comparison with uninjected | P value for comparison with Kesv 1 |
| --- | --- | --- | --- | --- |
| Kesv 1 | 15 | -33.47 ± 1.47 | <0.0001 (***) | - |
| Kesv 2 | 13 | -18.69 ± 2.65 | 0.03104 (*) | <0.0001 (***) |
| Kesv 1-N Δ35 | 6 | -11.33 ± 0.61 | 0.92402 (ns) | <0.0001 (***) |
| uninjected | 4 | -6.75 ± 2.25 | - | <0.0001 (***) |

Test indicated by ns for $p > 0.05$ , \* for $p < 0.05$ , \*\* for $p < 0.01$ and \*\*\* for $p < 0.001$ .

**Table S8:**
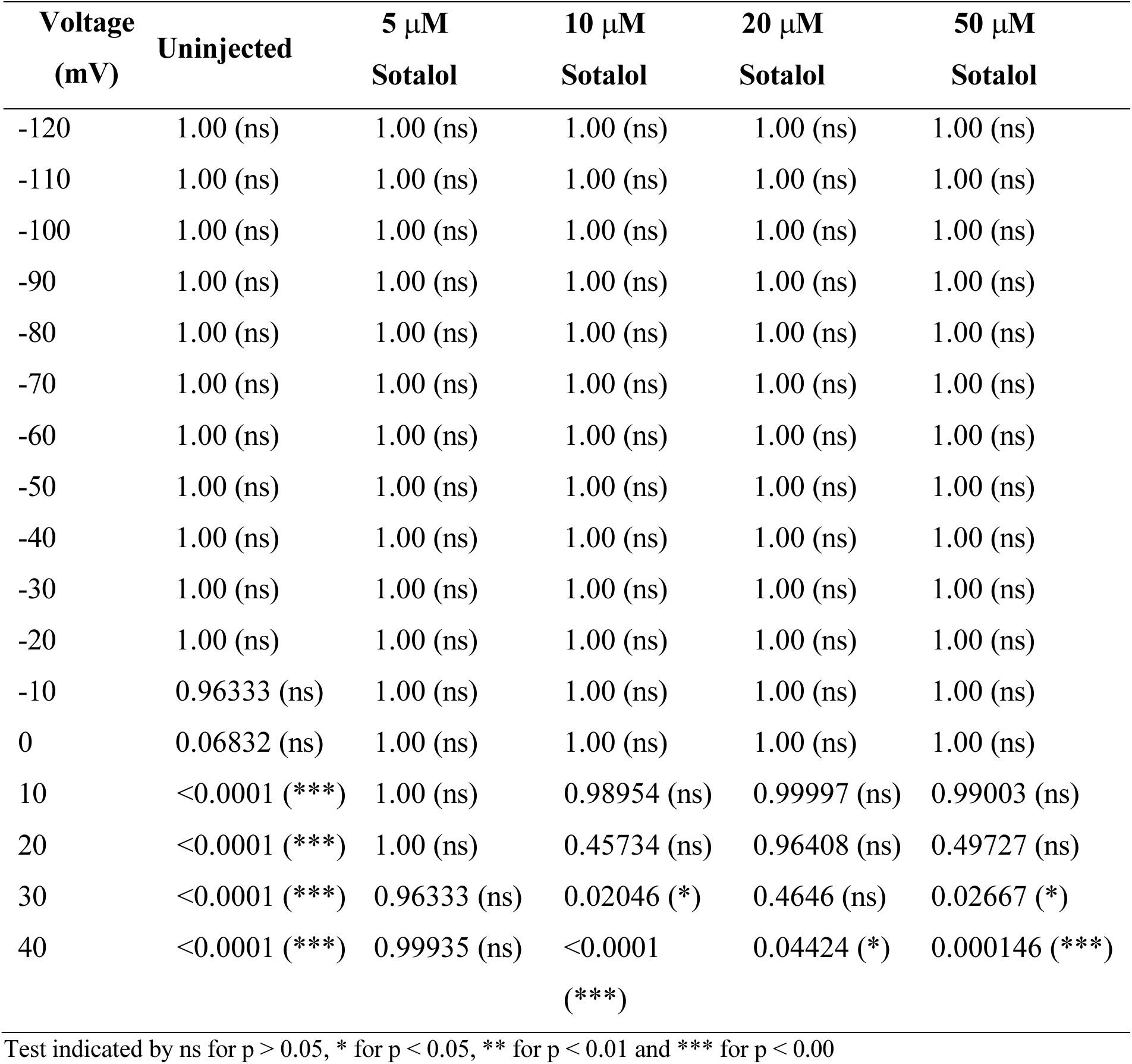
p-values for one-way ANOVA with posthoc mean comparison Tukey Test representing pharmacological activity of different concentrations of Sotalol on functional expression of Kesv 1.

| **Voltage**<br>**(mV)** | **Uninjected** | **5 μM**<br>**Sotalol** | **10 μM**<br>**Sotalol** | **20 μM**<br>**Sotalol** | **50 μM**<br>**Sotalol** |
| --- | --- | --- | --- | --- | --- |
| -120 | 1.00 (ns) | 1.00 (ns) | 1.00 (ns) | 1.00 (ns) | 1.00 (ns) |
| -110 | 1.00 (ns) | 1.00 (ns) | 1.00 (ns) | 1.00 (ns) | 1.00 (ns) |
| -100 | 1.00 (ns) | 1.00 (ns) | 1.00 (ns) | 1.00 (ns) | 1.00 (ns) |
| -90 | 1.00 (ns) | 1.00 (ns) | 1.00 (ns) | 1.00 (ns) | 1.00 (ns) |
| -80 | 1.00 (ns) | 1.00 (ns) | 1.00 (ns) | 1.00 (ns) | 1.00 (ns) |
| -70 | 1.00 (ns) | 1.00 (ns) | 1.00 (ns) | 1.00 (ns) | 1.00 (ns) |
| -60 | 1.00 (ns) | 1.00 (ns) | 1.00 (ns) | 1.00 (ns) | 1.00 (ns) |
| -50 | 1.00 (ns) | 1.00 (ns) | 1.00 (ns) | 1.00 (ns) | 1.00 (ns) |
| -40 | 1.00 (ns) | 1.00 (ns) | 1.00 (ns) | 1.00 (ns) | 1.00 (ns) |
| -30 | 1.00 (ns) | 1.00 (ns) | 1.00 (ns) | 1.00 (ns) | 1.00 (ns) |
| -20 | 1.00 (ns) | 1.00 (ns) | 1.00 (ns) | 1.00 (ns) | 1.00 (ns) |
| -10 | 0.96333 (ns) | 1.00 (ns) | 1.00 (ns) | 1.00 (ns) | 1.00 (ns) |
| 0 | 0.06832 (ns) | 1.00 (ns) | 1.00 (ns) | 1.00 (ns) | 1.00 (ns) |
| 10 | <0.0001 (***) | 1.00 (ns) | 0.98954 (ns) | 0.99997 (ns) | 0.99003 (ns) |
| 20 | <0.0001 (***) | 1.00 (ns) | 0.45734 (ns) | 0.96408 (ns) | 0.49727 (ns) |
| 30 | <0.0001 (***) | 0.96333 (ns) | 0.02046 (*) | 0.4646 (ns) | 0.02667 (*) |
| 40 | <0.0001 (***) | 0.99935 (ns) | <0.0001 (***) | 0.04424 (*) | 0.000146 (***) |

Test indicated by ns for $p > 0.05$ , \* for $p < 0.05$ , \*\* for $p < 0.01$ and \*\*\* for $p < 0.00$

**Table S9.** p-values for one-way ANOVA with posthoc mean comparison Tukey Test representing pharmacological activity of different concentrations of Linopirdine on functional expression of Kesv 1-DMSO.

| Voltage<br>(mV) | Uninjected-DMSO | 0.1 μM | 1 μM | 5 μM | 10 μM | 20 μM | 50 μM |
| --- | --- | --- | --- | --- | --- | --- | --- |
| -120 | 1.00 (ns) | 1.00 (ns) | 1.00 (ns) | 1.00 (ns) | 1.00 (ns) | 1.00 (ns) | 1.00 (ns) |
| -110 | 1.00 (ns) | 1.00 (ns) | 1.00 (ns) | 1.00 (ns) | 1.00 (ns) | 1.00 (ns) | 1.00 (ns) |
| -100 | 1.00 (ns) | 1.00 (ns) | 1.00 (ns) | 1.00 (ns) | 1.00 (ns) | 1.00 (ns) | 1.00 (ns) |
| -90 | 1.00 (ns) | 1.00 (ns) | 1.00 (ns) | 1.00 (ns) | 1.00 (ns) | 1.00 (ns) | 1.00 (ns) |
| -80 | 1.00 (ns) | 1.00 (ns) | 1.00 (ns) | 1.00 (ns) | 1.00 (ns) | 1.00 (ns) | 1.00 (ns) |
| -70 | 1.00 (ns) | 1.00 (ns) | 1.00 (ns) | 1.00 (ns) | 1.00 (ns) | 1.00 (ns) | 1.00 (ns) |
| -60 | 1.00 (ns) | 1.00 (ns) | 1.00 (ns) | 1.00 (ns) | 1.00 (ns) | 1.00 (ns) | 1.00 (ns) |
| -50 | 1.00 (ns) | 1.00 (ns) | 1.00 (ns) | 1.00 (ns) | 1.00 (ns) | 1.00 (ns) | 1.00 (ns) |
| -40 | 1.00 (ns) | 1.00 (ns) | 1.00 (ns) | 1.00 (ns) | 1.00 (ns) | 1.00 (ns) | 1.00 (ns) |
| -30 | 1.00 (ns) | 1.00 (ns) | 1.00 (ns) | 1.00 (ns) | 1.00 (ns) | 1.00 (ns) | 1.00 (ns) |
| -20 | 1.00 (ns) | 1.00 (ns) | 1.00 (ns) | 1.00 (ns) | 1.00 (ns) | 1.00 (ns) | 1.00 (ns) |
| -10 | 0.99872 (ns) | 1.00 (ns) | 1.00 (ns) | 1.00 (ns) | 1.00 (ns) | 1.00 (ns) | 1.00 (ns) |
| 0 | 0.39457 (ns) | 0.9963 (ns) | 1.00 (ns) | 1.00 (ns) | 1.00 (ns) | 1.00 (ns) | 1.00 (ns) |
| 10 | 0.00271 (**) | 0.4881 (ns) | 1.00 (ns) | 1.00 (ns) | 0.99801<br>(ns) | 1.00 (ns) | 1.00 (ns) |
| 20 | <0.0001<br>(***) | 0.01396 (*) | 1.00 (ns) | 0.87994<br>(ns) | 0.39572<br>(ns) | 0.99801<br>(ns) | 1.00 (ns) |
| 30 | <0.0001<br>(***) | <0.0001<br>(***) | 1.00 (ns) | 0.06499<br>(ns) | 0.00367<br>(**) | 0.88892<br>(ns) | 1.00 (ns) |
| 40 | <0.0001<br>(***) | <0.0001<br>(***) | 1.00 (ns) | 0.00015<br>(***) | <0.0001<br>(***) | 0.12895<br>(ns) | 1.00 (ns) |

Test indicated by ns for $p > 0.05$ , \* for $p < 0.05$ , \*\* for $p < 0.01$ and \*\*\* for $p < 0.001$

**Table S10.** p-values for one-way ANOVA with posthoc mean comparison Tukey Test representing pharmacological activity of different concentrations of Retigabine on functional expression of Kesv 1-DMSO.

| **Voltage**<br>**(mV)** | **Uninjected-**<br>**DMSO** | **5 μM** | **10 μM** | **20 μM** | **50 μM** |
| --- | --- | --- | --- | --- | --- |
| -120 | 1.00 (ns) | 1.00 (ns) | 1.00 (ns) | 1.00 (ns) | 1.00 (ns) |
| -110 | 1.00 (ns) | 1.00 (ns) | 1.00 (ns) | 1.00 (ns) | 1.00 (ns) |
| -100 | 1.00 (ns) | 1.00 (ns) | 1.00 (ns) | 1.00 (ns) | 1.00 (ns) |
| -90 | 1.00 (ns) | 1.00 (ns) | 1.00 (ns) | 1.00 (ns) | 1.00 (ns) |
| -80 | 1.00 (ns) | 1.00 (ns) | 1.00 (ns) | 1.00 (ns) | 1.00 (ns) |
| -70 | 1.00 (ns) | 1.00 (ns) | 1.00 (ns) | 1.00 (ns) | 1.00 (ns) |
| -60 | 1.00 (ns) | 1.00 (ns) | 1.00 (ns) | 1.00 (ns) | 1.00 (ns) |
| -50 | 1.00 (ns) | 1.00 (ns) | 1.00 (ns) | 1.00 (ns) | 1.00 (ns) |
| -40 | 1.00 (ns) | 1.00 (ns) | 1.00 (ns) | 1.00 (ns) | 1.00 (ns) |
| -30 | 1.00 (ns) | 1.00 (ns) | 1.00 (ns) | 1.00 (ns) | 1.00 (ns) |
| -20 | 1.00 (ns) | 1.00 (ns) | 1.00 (ns) | 1.00 (ns) | 1.00 (ns) |
| -10 | 1.00 (ns) | 1.00 (ns) | 1.00 (ns) | 1.00 (ns) | 1.00 (ns) |
| 0 | 1.00 (ns) | 1.00 (ns) | 1.00 (ns) | 1.00 (ns) | 1.00 (ns) |
| 10 | 0.97794 (ns) | 1.00 (ns) | 1.00 (ns) | 1.00 (ns) | 1.00 (ns) |
| 20 | 0.31087 (ns) | 1.00 (ns) | 1.00 (ns) | 1.00 (ns) | 1.00 (ns) |
| 30 | 0.00952 (***) | 1.00 (ns) | 1.00 (ns) | 1.00 (ns) | 0.99537 (ns) |
| 40 | <0.0001 (***) | 1.00 (ns) | 1.00 (ns) | 1.00 (ns) | 0.82192 (ns) |

Test indicated by ns for $p > 0.05$ , \* for $p < 0.05$ , \*\* for $p < 0.01$ and \*\*\* for $p < 0.001$

**Table S11.** Average distances between C*_α_* atoms of representative residues on opposite monomers in the MD simulations of Kesv 1 **(a)** and Kesv 2 **(b)** in comparison to average distances in PDB structures of open and closed conformations of MthK (PDB IDs 6U5R, 8DJB, 5BKI for closed and 1LNQ, 3LDC, 4QE9, 6OLY, 6U9P for open conformations) and KcsA (PDB IDs 1BL8, 1F6G, 5J9P, 2QTO, 3EFF, 7MHR for closed and 3F5W, 5VKE, 7M2J, 7MUB for open conformations).

| Kesv |  |  | MthK |  |  | KcsA |  |  |
| --- | --- | --- | --- | --- | --- | --- | --- | --- |
| residue | Kesv1 (nm) | Kesv2 (nm) | residue | closed (nm) | open (nm) | residue | closed (nm) | open (nm) |
| **S86** | 0.83 | 0.83 | **T59** | 0.86 | 0.83 | **T75** | 0.91 | 0.89 |
| **L115** | 2.14 | 1.97 | **A88** | 1.87 | 1.55 | **G104** | 1.18 | 1.56 |
| **L38** | 2.62 | 2.61 | **P19** | -- | 3.74 | **A31** | 3.72 | 3.86 |
| **K124** | 1.18 | 1.30 | **F97** | 2.06 | 3.60 | **T112** | 1.17 | 2.46 |

**Table S12:**
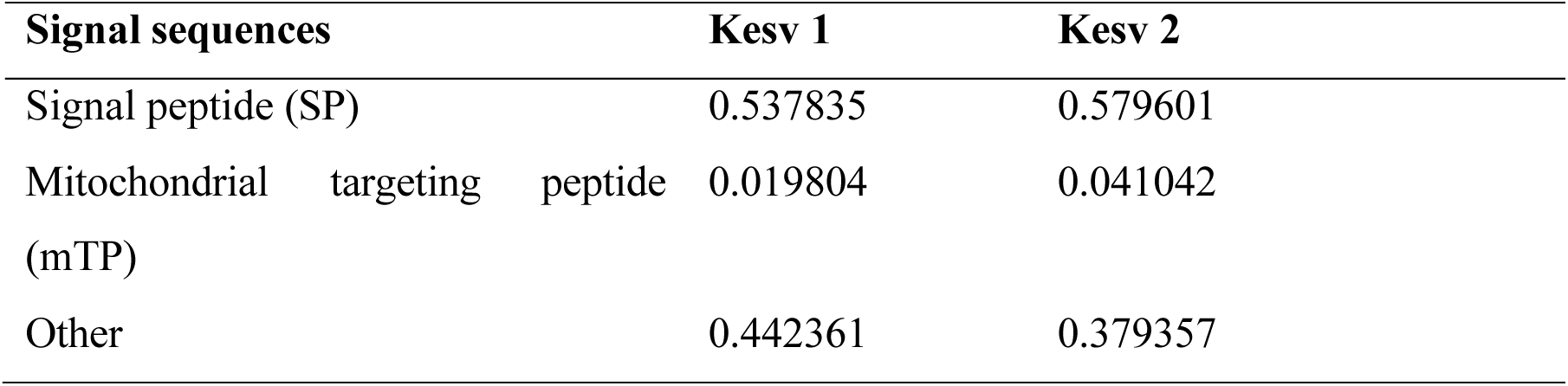
Probability values of N-terminal presequences for subcellular location of Kesv 1 and Kesv 2 based on TargetP-2.0 tool.

| Signal sequences | Kesv 1 | Kesv 2 |
| --- | --- | --- |
| Signal peptide (SP) | 0.537835 | 0.579601 |
| Mitochondrial targeting peptide (mTP) | 0.019804 | 0.041042 |
| Other | 0.442361 | 0.379357 |

